# ESX-1 substrates ESAT-6 and CFP-10 limit cGAS-STING activation via DNA and Zn^2+^ restriction

**DOI:** 10.64898/2026.10.26.761845

**Authors:** Lei Xu, Jianan Huang, Damin Pan, Conghui Wu, Tianyuan Chang, Jiaxing Lu, Xinfang Zhao, Hongjin Wang, Shuyu Li, Lingfang Zhu, Yicheng Sun, Yao Wang, Xihui Shen

**Affiliations:** State Key Laboratory for Crop Stress Resistance and High-Efficiency Production, Shaanxi Key Laboratory of Agricultural and Environmental Microbiology, College of Life Sciences, Northwest A&F University, Yangling, Shaanxi 712100, China; Department of Biochemistry and Molecular Biology, College of Life Sciences, Northwest A&F University, Yangling 712100, China; NHC Key Laboratory of Systems Biology of Pathogens, State Key Laboratory of Respiratory Health and Multimorbidity, National Institute of Pathogen Biology and Center for Tuberculosis Research, Chinese Academy of Medical Sciences & Peking Union Medical College, Beijing 102629, China

**Keywords:** *Mycobacterium tuberculosis*, ESX-1, cGAS-STING, ESAT-6, CFP-10, cytosolic DNA sensing, zinc restriction, immune evasion

## Abstract

ESX-1–mediated phagosomal permeabilization exposes mycobacterial DNA to cytosolic cGAS–STING surveillance, yet the response is not overwhelming, suggesting additional layers of control exist. Here we show that the ESX-1 substrates ESAT-6 and CFP-10 constrain cGAS activation through complementary biochemical mechanisms. Purified *Mycobacterium tuberculosis* CFP-10 is a metal-dependent, 5′–3′ exonuclease-like enzyme that preferentially degrades linear DNA, limiting the availability of stimulatory ligand. By contrast, purified *M. tuberculosis* ESAT-6 binds Zn^2+^ and limits the Zn^2+^-dependent conditions required for cGAS activation. During *M. marinum* infection, loss of these substrates augmented interferon-associated responses. Wild-type alleles restored immune suppression, whereas DNA-processing-defective CFP-10 or Zn^2+^-binding-defective ESAT-6 variants did not. These genotype-dependent differences were largely abolished in *cGas^−/–^* and *Sting^−/–^* macrophages. The double mutant was attenuated in zebrafish and mice, and its enhanced immune activation and attenuation were markedly diminished in *cGas^−/–^* mice. Prior exposure to the mutant improved zebrafish survival following wild-type challenge. Thus, beyond enabling mycobacterial DNA access to the cytosol, ESX-1 deploys its substrates to constrain cGAS–STING signaling by controlling activating-ligand availability and the metal-dependent environment required for sensor activation.

**Graphical abstract:** 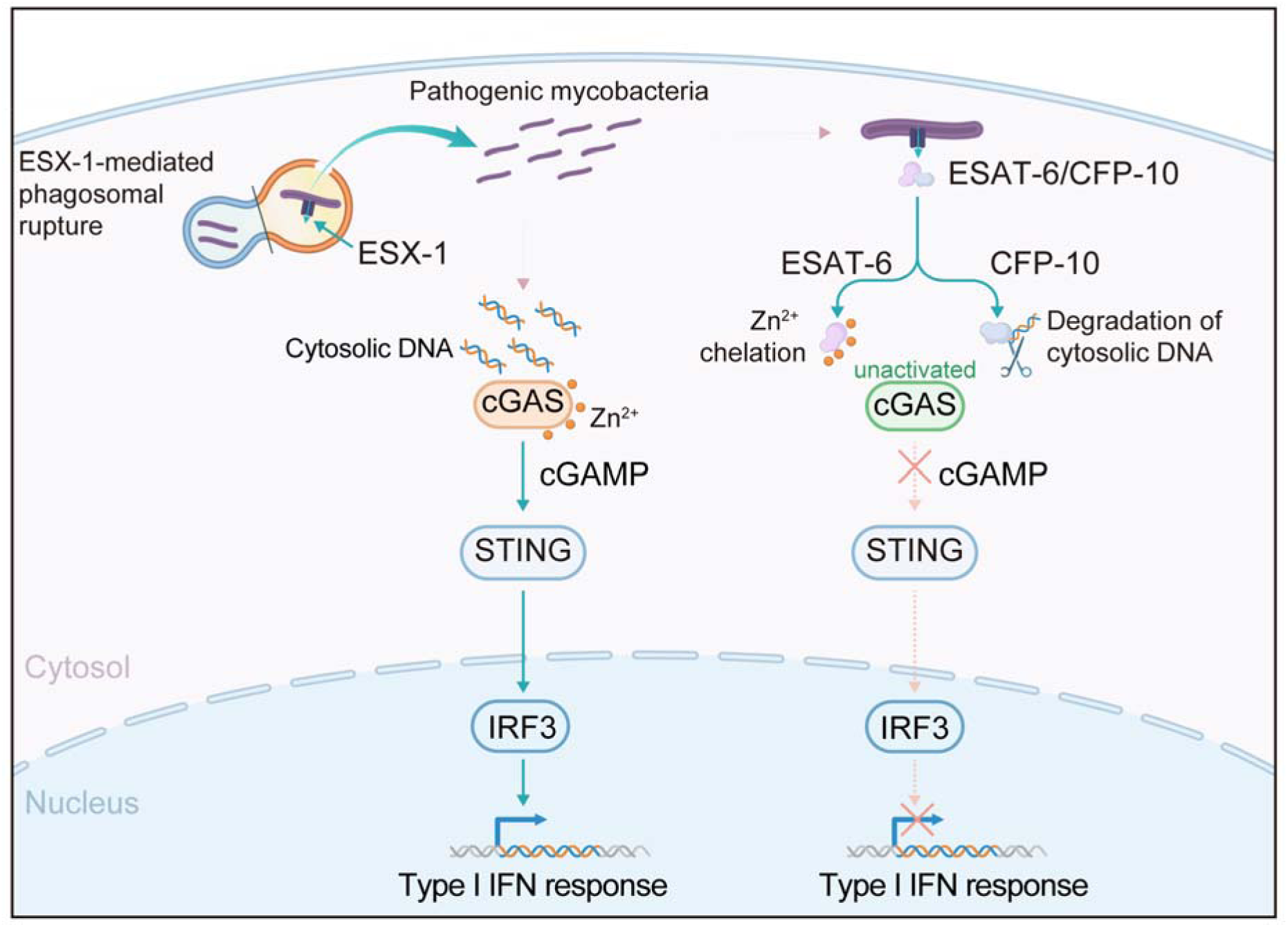

**Highlights:**

- ESAT-6 and CFP-10 possess complementary activities that limit cGAS-STING activation
- CFP-10 acts as a 5′-3′ exonuclease that degrades linear immunostimulatory DNA
- ESAT-6 binds Zn^2+^ and restricts the ionic conditions required for cGAS activation
- Dual disruption of ESAT-6/CFP-10 attenuates virulence and enhances protective priming

**IMPORTANCE:** ESX-1 is best known for breaching the mycobacterial phagosome, thereby exposing DNA to the cytosolic sensor cGAS-STING. We show that two major ESX-1 substrates also tune what happens after cytosolic access. CFP-10 reduces the availability of stimulatory DNA, whereas ESAT-6 limits the Zn^2+^-dependent conditions needed for efficient cGAS activation. These complementary activities restrain cGAS-dependent immunity and promote mycobacterial virulence *in vivo*. The findings add a regulatory layer to, rather than replace, the established ESX-1 model and reveal a broader principle of immune evasion: pathogens can suppress innate sensors by controlling both their activating ligands and the physicochemical environment required for signaling.

## INTRODUCTION

*Mycobacterium tuberculosis* (*Mtb*) remains a leading cause of human mortality, and its success as an intracellular pathogen depends critically on the ESX-1 type VII secretion system. ESX-1 permeabilizes the phagosomal membrane, delivers bacterial products into the host cytosol, and is essential for full virulence^1–4^. A direct consequence of this cytosolic access is the exposure of mycobacterial and host-derived DNA to innate immune surveillance^5–7^.

The cyclic GMP-AMP synthase (cGAS)–STING pathway is the principal sensor of cytosolic DNA. Upon binding double-stranded DNA, cGAS synthesizes 2′3′-cGAMP, which activates STING and downstream TBK1–IRF3 signaling ^8^. During mycobacterial infection, cGAS detects *M. tuberculosis* DNA and drives type I interferon production^1,9,10^, establishing ESX-1-mediated membrane damage as a key trigger of cytosolic DNA sensing. Yet this conventional view collides with a long-standing paradox: highly virulent, ESX-1-intact mycobacteria often elicit weaker innate responses than attenuated or ESX-1-compromised strains, including reduced type I interferon induction {Shah, 2013 #481}{Wassermann, 2015 #10}. Pathogenic mycobacteria thus appear to use ESX-1 not only to open the phagosome but also to restrain responses that would otherwise compromise bacterial survival. The intensity of cGAS–STING activation is therefore likely under active selection, particularly because type I interferon signaling can be host-protective or host-detrimental in tuberculosis depending on context ^11–13^.

Productive cGAS signaling requires more than the presence of cytosolic DNA. DNA binding drives higher-order cGAS assembly and condensation, and Zn^2+^ promotes DNA-induced cGAS condensation and catalytic activity ^8,14^. cGAS can therefore be regulated at two genetically separable levels: the availability and physical state of stimulatory DNA, and the physicochemical conditions that permit productive sensor assembly. Whether mycobacterial effectors exploit either control point has not been established. This gap matters because the ESX-1 system exports not only pore-forming effectors but also abundant substrates whose fate and function after secretion remain poorly defined. ESAT-6 and CFP-10 are the best-characterized ESX-1 substrates and form a stable heterodimer ^15^. ESAT-6 has been linked to membrane interaction, phagosomal damage, and cytosolic translocation^16–18^, whereas CFP-10 is integrated into the broader ESX-1 secretion network^19–21^. These functions explain how ESAT-6 and CFP-10 contribute to ESX-1-dependent virulence, but they do not address what these proteins may do after secretion. Whether either substrate has additional biochemical activity capable of directly modulating host signaling remains largely unexplored. More generally, whether bacterial effectors can shape innate immune output by acting on the physicochemical requirements of a sensor rather than on the sensor or its ligand remains an open question.

To test this possibility, we combined purified-protein biochemistry with bacterial genetics, transcriptomics, cell-based assays, and in vivo infection, using *Mycobacterium marinum* as a genetically tractable, ESX-1-competent surrogate whose RD1/ESX-1 locus is syntenic with that of *Mtb* and whose ESAT-6 and CFP-10 are highly conserved ^15–17,20,22–24^. We show that ESAT-6 and CFP-10 possess complementary biochemical activities that limit DNA-triggered cGAS activation. CFP-10 is a metal-dependent 5′–3′ exonuclease that preferentially degrades linear immunostimulatory DNA, reducing ligand availability, whereas ESAT-6 binds Zn^2+^ and restricts the Zn^2+^-dependent conditions required for efficient cGAS assembly. Activity-defective alleles and cGAS–STING-deficient cells and mice link these activities to cGAS–STING suppression and virulence *in vivo*. These findings expand the established model of ESX-1 biology: ESX-1 promotes cytosolic access and DNA sensing, and individual secreted substrates additionally tune response intensity through complementary control of the activating ligand and its biochemical environment.

## RESULTS

### ESAT-6 and CFP-10 cooperatively suppress IFN-associated responses during infection

The ESX-1 secretion system is functionally conserved between *M. marinum* and *M. tuberculosis*, and *M. marinum* retains the major ESX-1 substrates ESAT-6 and CFP-10 while offering greater experimental tractability and lower pathogenicity ^21,25–27^. We used *M. marinum* as a surrogate model to investigate how ESAT-6 and CFP-10 regulate host innate immunity at both its optimal growth temperature (30°C) and a mammalian host-relevant temperature (37°C) ^26^. Infection at either temperature elicited rapid innate immune activation in primary macrophages (Figure S1A and S1B), with transcriptomic analysis revealing enrichment of NF-κB, Toll-like receptor, JAK–STAT, and cytosolic DNA-sensing pathways (Figure S1C). These findings establish *M. marinum* as a suitable model for dissecting ESAT-6/CFP-10-mediated immune modulation under both permissive and host-relevant conditions.

To define the contribution of ESAT-6 and CFP-10, we generated isogenic *M. marinum* deletion strains lacking *esat-6* (Δ*esat-6*), *cfp-10* (Δ*cfp-10*), or both (Δ*esat-6*Δ*cfp-10*). Comparative transcriptomic profiling of infected macrophages revealed that the double mutant induced a marked shift in host gene expression relative to wild-type (WT) infection, with numerous upregulated transcripts mapping to interferon (IFN)- and inflammation-associated programs (Figure 1A). Importantly, the deletion did not impair bacterial growth kinetics or early intracellular uptake (Figure S2A and S2B), ruling out indirect effects secondary to altered bacterial fitness or infection efficiency. Heatmap visualization showed progressively increased expression of *Ifnb1* and multiple interferon-stimulated genes (ISGs) in cells infected with single mutants, with the strongest response consistently observed in the Δ*esat-6*Δ*cfp-10* group (Figure 1B). Gene set enrichment analysis confirmed significant enrichment of the IFN-α response signature in Δ*esat-6*Δ*cfp-10*-infected macrophages compared with WT-infected cells (Figure 1C), indicating that simultaneous loss of ESAT-6 and CFP-10 unleashes a heightened type I IFN program. Furthermore, comparison of Δ*esat-6*Δ*cfp-10*-infected macrophages with mock-treated cells showed robust transcriptional induction of inflammatory and IFN-associated genes (Figure S2C). In addition, the IFN-γ response signature was significantly enriched in Δ*esat-6*Δ*cfp-10*-infected macrophages relative to WT-infected cells (Figure S2D), suggesting that loss of ESAT-6 and CFP-10 broadly amplifies IFN-associated inflammatory programs.

**Figure 1.**
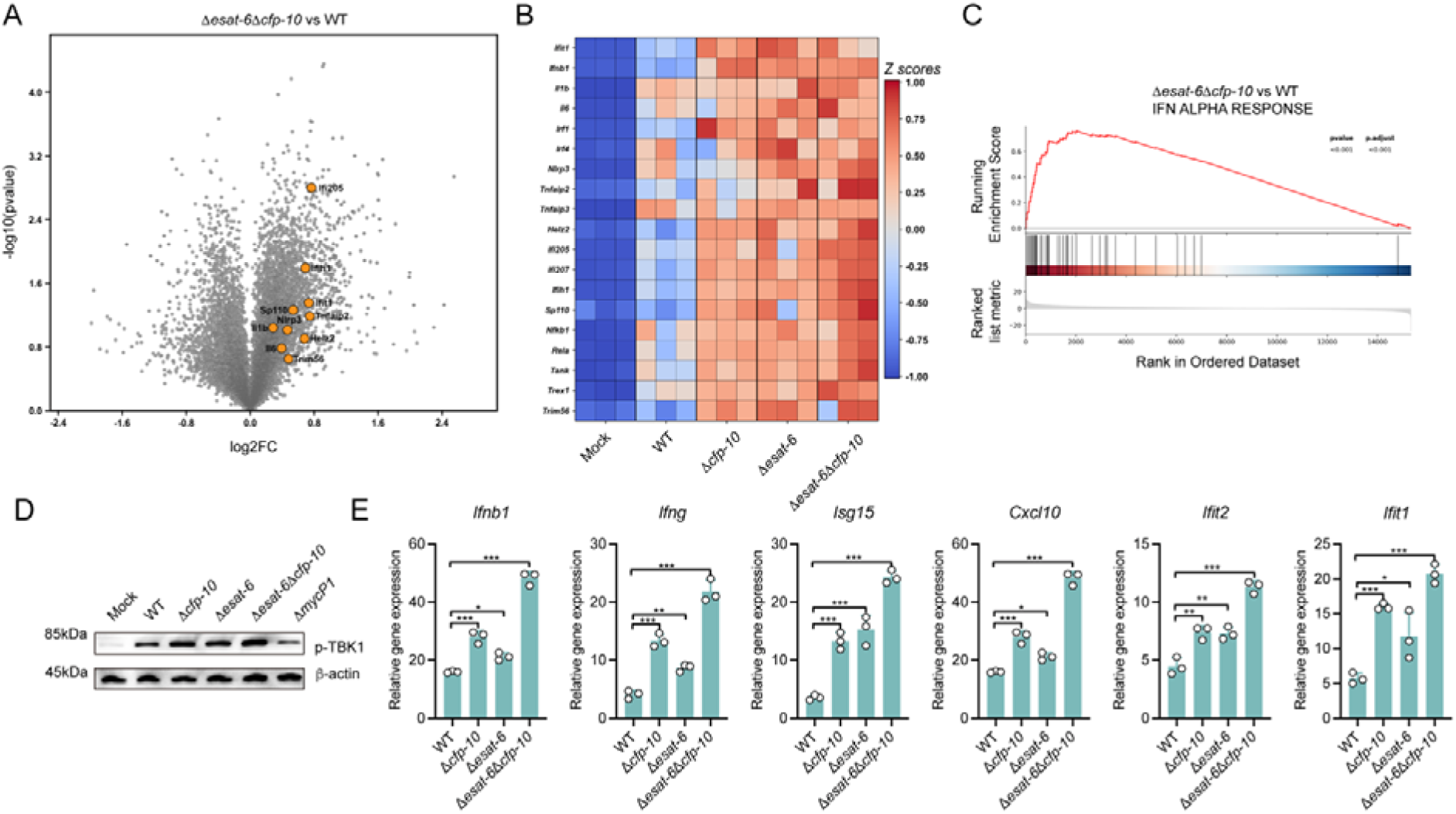
ESAT-6 and CFP-10 restrain interferon-associated responses in macrophages during mycobacterial infection. (A) Volcano plot showing differentially expressed genes identified by RNA-seq in primary macrophages infected with the Δ*esat-6*Δ*cfp-10* strain (MOI = 20, 4 h) relative to WT *M. marinum*. n = 3. (B) Heatmap showing Z-score-normalized expression of selected immune-related genes across the indicated conditions (Mock, WT, Δ*esat-6*, Δ*cfp-10* and Δ*esat-6*Δ*cfp-10*) (MOI = 20, 4 h) from the RNA-seq dataset. n = 3. (C) Gene set enrichment analysis (GSEA) performed on ranked transcriptomic data comparing Δ*esat-6*Δ*cfp-10*-infected macrophages with WT-infected macrophages using the IFN ALPHA RESPONSE gene set. n = 3. (D) Immunoblot analysis of phosphorylated TBK1 (p-TBK1) in primary macrophages infected with the indicated strains (MOI = 20, 4 h). β-actin was used as a loading control. (E) RT-qPCR analysis of indicated genes in primary macrophages infected with WT, Δ*cfp-10*, Δ*esat-6*, or Δ*esat-6*Δ*cfp-10* strains (MOI = 20, 4 h). Gene expression was normalized to the indicated internal control and presented as relative expression values. n = 3. *P* values were calculated using two-tailed Student’s t-test for paired comparisons or one-way or two-way ANOVA for multiple comparisons. Error bars represent ± SEM. \**P* < 0.05; \*\**P* < 0.01; \*\*\**P* < 0.001; ns, not significant.

Infection with mutant strains led to stronger activation of the upstream pathway, as evidenced by increased TBK1 phosphorylation relative to WT infection (Figure 1D). Quantitative PCR in primary macrophages validated the transcriptomic findings, with deletion of *esat-6* or *cfp-10* significantly increasing expression of *Ifnb1*, *Ifng*, and many ISGs, with the double mutant consistently eliciting the highest expression levels (Figure 1E). The enhanced expression of IFN-associated genes in response to mutant strains was also reproduced in RAW 264.7 macrophages, again with the strongest response in the double-mutant group (Figure S2E). These data indicate that ESAT-6 and CFP-10 cooperatively suppress IFN-associated responses in macrophages during early mycobacterial infection, prompting us to define their direct biochemical functions.

### CFP-10 is a metal-dependent 5′–3′ exonuclease that targets linear DNA

*In silico* analysis predicted a potential nucleic acid-interacting surface in CFP-10, with candidate DNA-contacting residues clustered on the protein surface (Figure 2A). We purified recombinant CFP-10 (Figure S3A and S3B), and electrophoretic mobility shift assays (EMSA) using short linear dsDNA substrates did not yield a stable retarded complex. Instead, the free DNA signal progressively decreased as CFP-10 input increased, whereas BSA had no such effect (Figure 2B), and isothermal titration calorimetry (ITC) confirmed direct binding of CFP-10 to DNA (Figure 2C). This pattern suggested that CFP-10 does not simply bind DNA stably but rather processes the substrate.

**Figure 2.**
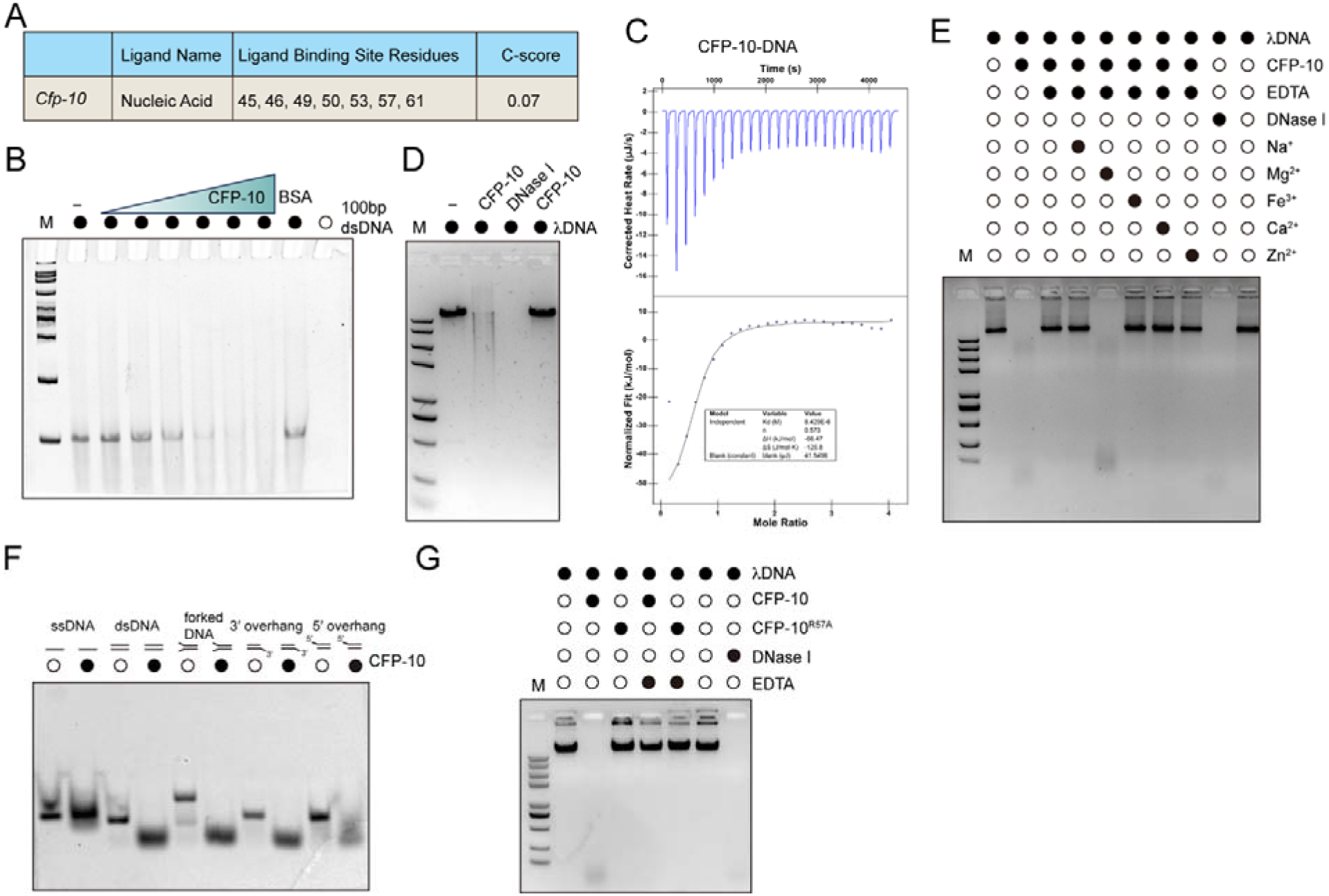
CFP-10 functions as a metal-dependent DNase. (A) *In silico* prediction of a putative nucleic acid-binding surface in CFP-10, showing the predicted ligand type, candidate binding residues, and confidence score. Putative binding surfaces were inferred from structural modeling (https://alphafold.com/) and molecular docking analyses (https://aideepmed.com/). (B) Electrophoretic mobility shift assay (EMSA) using a 100-bp dsDNA (40 ng) substrate incubated with increasing amounts of recombinant *M. tuberculosis* CFP-10 (1 μM, 3 μM, 6 μM, 9 μM, 12 μM, and 15 μM) or BSA (8 μM), followed by native gel electrophoresis. (C) Isothermal titration calorimetry (ITC) analysis of recombinant *M. tuberculosis* CFP-10 (25 μM) with 100-bp dsDNA (500 μM). Upper panel, raw heat change upon sequential injections; lower panel, fitted binding isotherm. (D) *In vitro* DNA degradation assay in which linear λDNA (20 ng) was incubated with recombinant *M. tuberculosis* CFP-10 (10 μM), DNase I (1 U/μL), or heat-inactivated CFP-10 (10 μM) and analyzed by agarose gel electrophoresis. (E) Metal-dependence assay in which λDNA (20 ng) was incubated with recombinant *M. tuberculosis* CFP-10 (10 μM) in the presence of EDTA (25 mM) or the indicated metal ions (4 mM Na^⁺^, 4 mM Mg^2+^, 1 mM Fe^3+^, 4 mM Ca^2+^, and 1 mM Zn^2+^), followed by agarose gel analysis. (F) Substrate preference assay using DNA substrates with distinct terminal structures, including ssDNA (2 μM), blunt-ended dsDNA (2 μM), forked DNA (2 μM), 3′-overhang DNA (2 μM), and 5′-overhang DNA (2 μM), incubated with recombinant *M. tuberculosis* CFP-10 (10 μM) and analyzed by gel electrophoresis. (G) DNA degradation assay comparing λDNA (20 ng) processing by wild-type *M. tuberculosis* CFP-10 (10 μM) and the R57A mutant (10 μM) in the presence or absence of EDTA (25 mM).

Consistent with this hypothesis, incubation of purified *M. tuberculosis* CFP-10 with linear λDNA resulted in marked substrate degradation, whereas heat-inactivated protein lost this activity (Figure 2D). Comparable activity was observed with the *M. marinum* ortholog, indicating functional conservation (Figure S3C). Circular plasmid DNA remained largely resistant to CFP-10 treatment across the same concentration range, whereas CFP-10 efficiently processed linear DNA (Figures S3D and S3E), revealing a strong preference for linear substrates. Ectopic expression of wild-type or mutant *cfp-10* did not cause a major growth defect in bacterial expression systems, arguing against indiscriminate nuclease toxicity (Figure S3F).

Biochemically, DNA degradation by CFP-10 was strongly inhibited by EDTA, indicating a requirement for divalent metal ions (Figure 2E). Mg^2+^ restored robust activity, whereas other tested ions showed weaker or no support (Figure 2E; Figure S3G). To further delineate its mode of action, we utilized linear dsDNA substrates designed with distinct terminal structures, including 5′ overhangs, 3′ overhangs, and substrates with both ends protruding. CFP-10 exhibited a clear preference for substrates compatible with 5′-to-3′ processing, demonstrating that it functions primarily as a 5′–3′ exonuclease rather than an indiscriminate endonuclease (Figure 2F).

Further analysis nominated residue R57 as a candidate DNA-contact site within a highly conserved region (Figure S3H and S3I). Substitution of R57 with alanine markedly impaired DNA degradation by both *M. tuberculosis* and *M. marinum* CFP-10 *in vitro* (Figure 2G; Figure S3J). Finally, purified CFP-10 efficiently degraded mycobacterial genomic DNA *in vitro*, whereas the R57A mutant and heat-inactivated protein were largely inactive (Figure S3K–S3M). Together, these data identify CFP-10 as a metal-dependent exonuclease-like factor that preferentially processes linear DNA.

### CFP-10 limits cGAS activation by depleting stimulatory DNA

Having established that CFP-10 possesses exonuclease-like activity, we next asked whether this activity is sufficient to restrain DNA-dependent cGAS activation. Using an HPLC-based *in vitro* cGAS reconstitution system, addition of recombinant cGAS to ATP/GTP and stimulatory dsDNA led to robust 2′3′-cGAMP production (Figure S4A). Under these conditions, addition of recombinant CFP-10 markedly reduced cGAMP production in a concentration-dependent manner (Figure 3A). This effect depended on the DNA-processing activity of CFP-10, as both heat-inactivated protein and the DNA-processing-defective CFP-10^R57A^ mutant showed substantially reduced inhibitory activity (Figure 3B).

**Figure 3.**
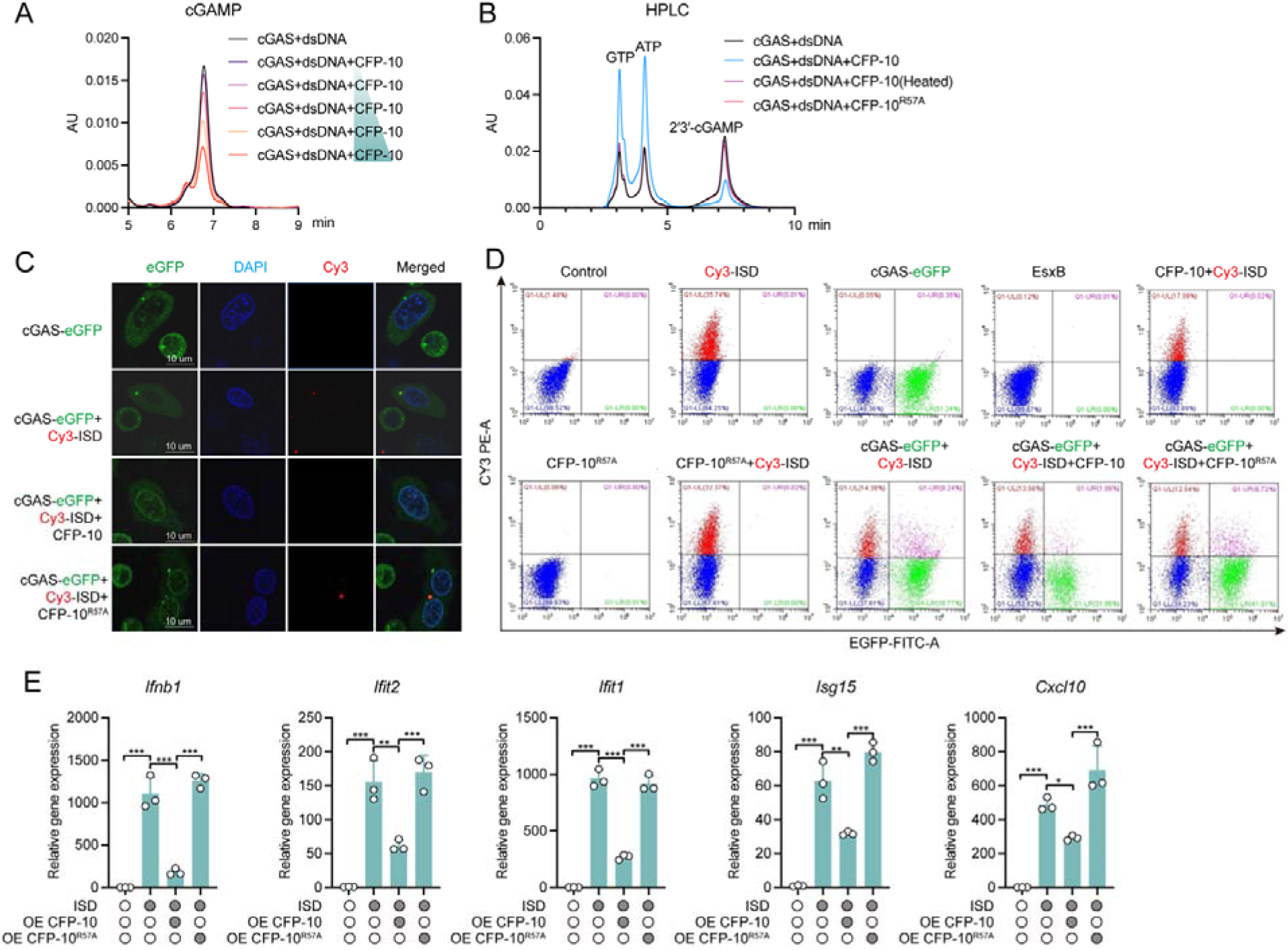
CFP-10 suppresses DNA-triggered cGAS activation in biochemical and cellular assays. (A) HPLC analysis of 2′3′-cGAMP production in a reconstituted in vitro cGAS assay containing recombinant cGAS (10 μM), dsDNA (10^-2^ mg/mL), ATP (0.2 mM), and GTP (0.2 mM), with increasing amounts of recombinant CFP-10 (0, 5, 10, 15, 20 μM) added as indicated. (B) HPLC analysis of nucleotide products from the *in vitro* cGAS reaction performed in the presence of wild-type CFP-10 (10 μM), heat-inactivated CFP-10 (10 μM), or CFP-10^R57A^ (10 μM). Peaks corresponding to GTP, ATP, and 2′3′-cGAMP are indicated. (C) Confocal microscopy of HeLa cells expressing cGAS-eGFP and transfected with Cy3-labeled ISD in the presence or absence of wild-type CFP-10 or CFP-10(R57A). Nuclei were stained with DAPI. Representative images from 3 independent experiments. (D) Flow cytometric analysis of HeLa cells expressing cGAS-eGFP and/or Cy3-labeled ISD together with WT CFP-10 or CFP-10^R57A^, as indicated. Representative plots from 3 independent experiments. (E) RT-qPCR analysis of gene expression in cells transfected with ISD and expressing wild-type CFP-10 or CFP-10^R57A^. Gene expression was normalized to the indicated internal control and presented as relative expression values. *P* values were calculated using two-tailed Student’s t-test for paired comparisons or one-way or two-way ANOVA for multiple comparisons. Error bars represent ± SEM. \**P* < 0.05; \*\**P* < 0.01; \*\*\**P* < 0.001; ns, not significant.

We next tested whether this biochemical effect could be recapitulated in cells. In HeLa cells expressing cGAS-eGFP, transfection of Cy3-labeled ISD induced prominent cGAS puncta, consistent with DNA-triggered higher-order cGAS assembly. Co-expression of wild-type CFP-10 markedly reduced cGAS puncta formation, whereas CFP-10^R57A^ exhibited a much weaker effect (Figure 3C). Flow cytometry further showed that CFP-10 reduced the intracellular DNA-associated Cy3 signal, while the R57A mutant was less effective (Figure 3D), consistent with depletion of stimulatory DNA in cells. Functionally, wild-type CFP-10 significantly attenuated ISD-induced expression of *Ifnb1* and ISGs, whereas the CFP-10^R57A^ mutant displayed markedly impaired suppressive activity (Figures 3E and S4B). Together, these results support a model in which CFP-10 suppresses host DNA sensing by intercepting and processing stimulatory DNA before productive cGAS engagement.

### ESAT-6 binds Zn^2+^ through a conserved H26/D30 site

Because deletion of ESAT-6 also enhanced IFN-associated responses, we asked whether ESAT-6 restrains innate immunity through a mechanism distinct from that of CFP-10. *In silico* analysis predicted a potential Zn^2+^-binding site in ESAT-6, with residues H26 and D30 emerging as candidate coordinating residues (Figure 4A). ITC revealed that recombinant ESAT-6 bound Zn^2+^ with clear and saturable kinetics, yielding a dissociation constant in the submicromolar range, whereas binding to Ca^2+^, Cu^2+^, and Fe³^⁺^ was markedly weaker (Figure 4B; Figure S5B). In a PAR-based competition assay, formation of the PAR-Zn^2+^ complex generated a characteristic absorbance peak near 500 nm, and addition of recombinant ESAT-6 reduced this signal in a protein dose-dependent manner, consistent with direct competition for Zn^2+^ binding (Figure 4C). Heat-inactivated ESAT-6 largely lost this activity (Figure S5C).

**Figure 4.**
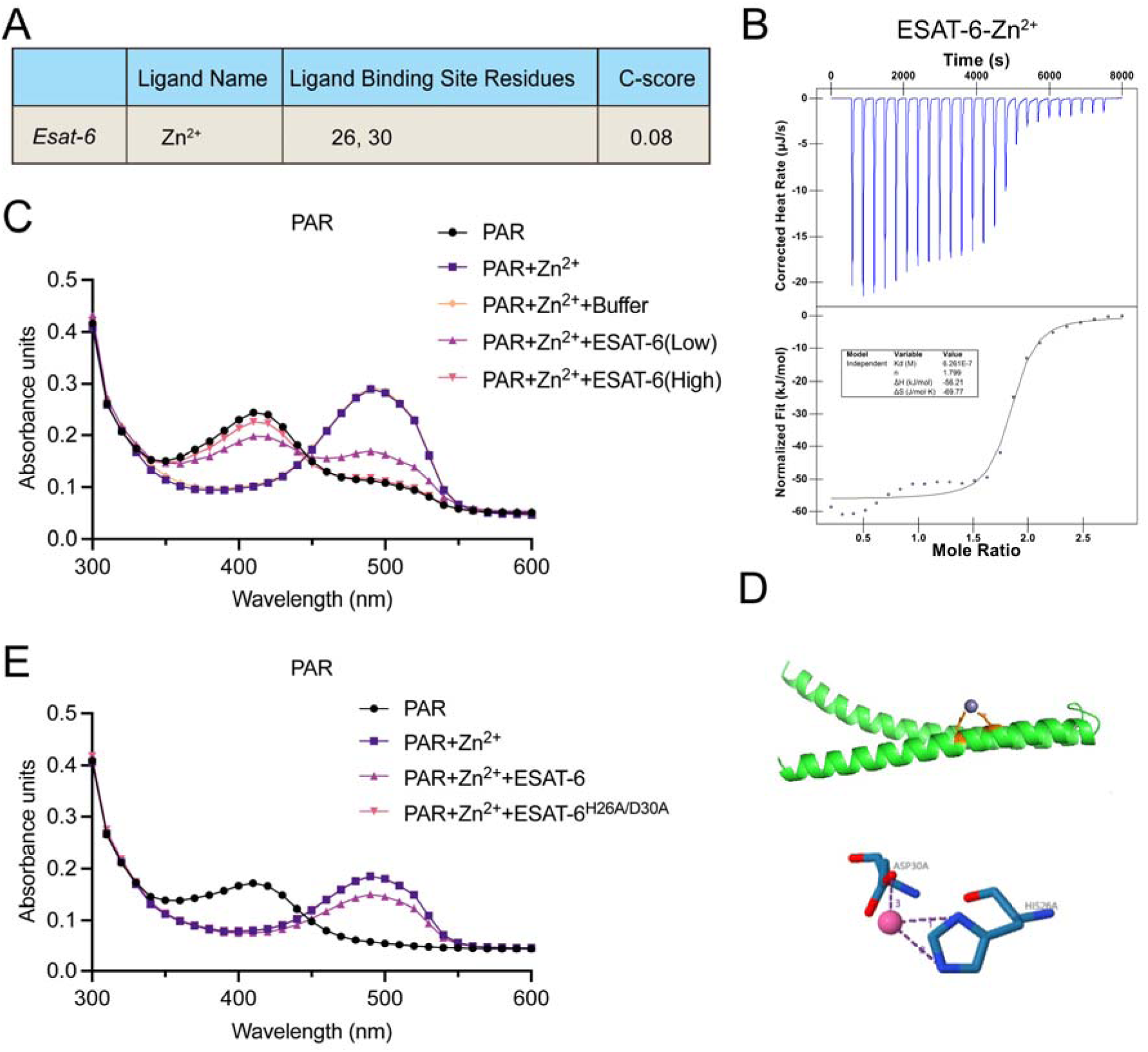
ESAT-6 is a Zn^2+^-binding protein. (A) *In silico* prediction of a putative Zn^2+^-binding site in ESAT-6, showing the predicted ligand type, candidate binding residues, and confidence score. Putative binding surfaces were inferred from structural modeling (https://alphafold.com/) and molecular docking analyses (https://aideepmed.com/). (B) Isothermal titration calorimetry (ITC) analysis of recombinant ESAT-6 (25 μM) with Zn^2+^ (1 mM). Upper panel, raw heat change upon sequential injections; lower panel, fitted binding isotherm. (C) PAR competition assay performed with PAR (10 μM), Zn^2+^ (5 μM), and increasing amounts of recombinant ESAT-6 (0.4, 0.67 μM), with absorbance spectra recorded across the indicated wavelengths. (D) Molecular docking model of Zn^2+^ bound to ESAT-6, highlighting the predicted coordinating residues H26 and D30. (E) PAR competition assay comparing WT ESAT-6 (0.4 μM) and the ESAT-6^H26A/D30A^ (2.0 μM) mutant for Zn^2+^ (5 μM) binding.

Molecular docking positioned Zn^2+^ within a discrete pocket of ESAT-6 and predicted direct coordination by H26 and D30 (Figure 4D). Multiple-sequence alignment showed that these residues are highly conserved among ESAT-6 homologs, and phylogenetic analysis supported broad conservation of this protein family (Figure S5D and S5E). The double mutant ESAT-6^H26A/D30A^ was markedly impaired in its ability to compete for Zn^2+^ relative to wild-type ESAT-6 in the PAR competition assay (Figure 4E), demonstrating that H26 and D30 are critical determinants of Zn^2+^ binding. Thus, ESAT-6 is a *bona fide* Zn^2+^-binding protein with a structurally defined coordination mechanism.

### ESAT-6 restrains cGAS activation by limiting Zn^2+^ availability

Previous work has shown that Zn^2+^ promotes cGAS activation by facilitating DNA-induced cGAS phase separation and enhancing its enzymatic activity ^14^. We next asked whether the Zn^2+^-binding activity of ESAT-6 is sufficient to restrain DNA-triggered cGAS activation. In the HPLC-based reconstituted cGAS assay, recombinant ESAT-6 reduced dsDNA-induced 2′3′-cGAMP production in a dose-dependent manner (Figure 5A). Importantly, the addition of exogenous Zn^2+^ restored cGAMP production under the same conditions, indicating that ESAT-6 suppresses cGAS activation by limiting Zn^2+^ availability rather than by directly blocking catalysis (Figure 5A). Consistent with this interpretation, both heat-inactivated ESAT-6 and the Zn^2+^-binding-defective mutant ESAT-6^H26A/D30A^ showed markedly reduced inhibitory activity (Figure 5B).

**Figure 5.**
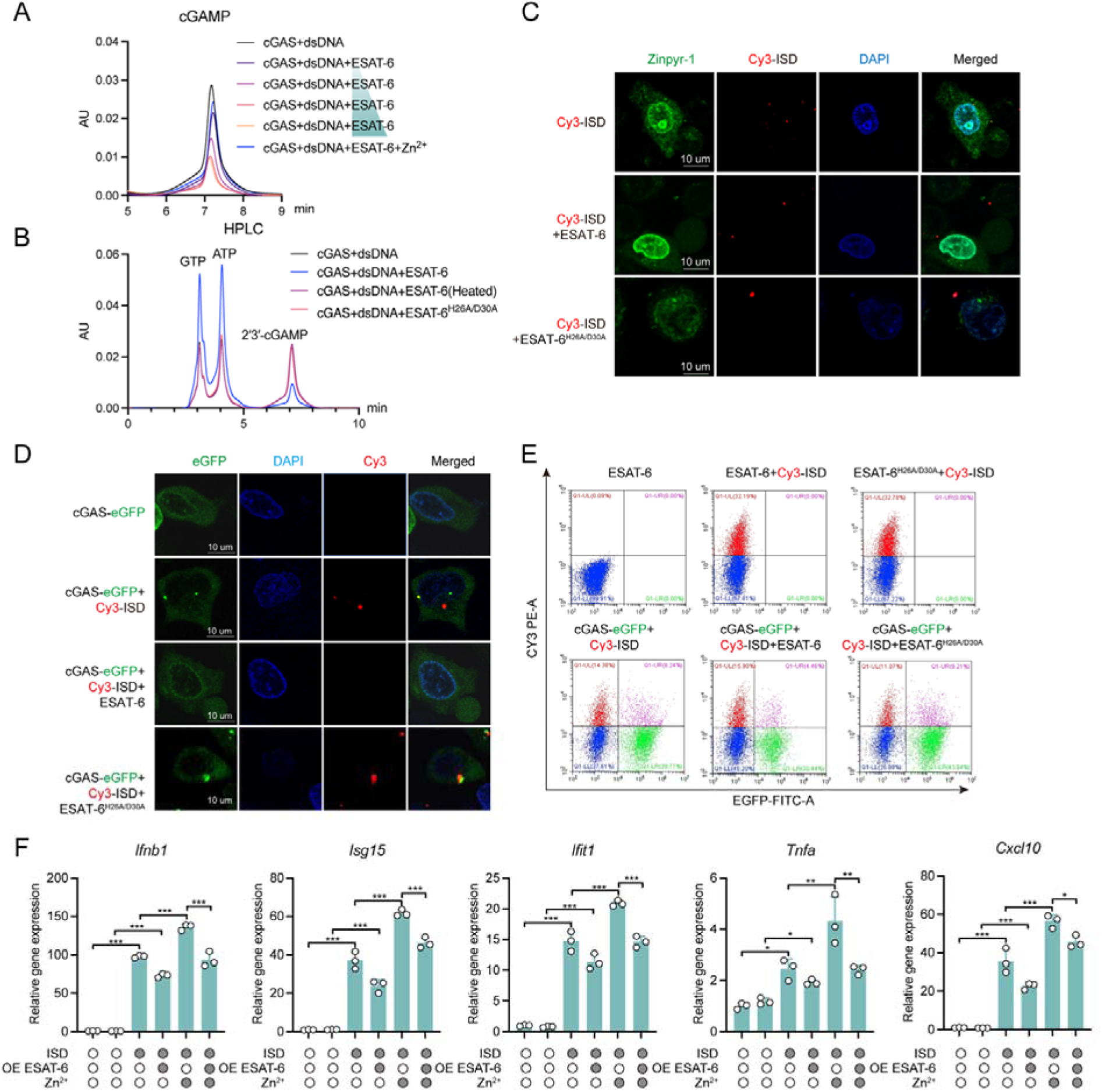
ESAT-6 suppresses DNA-triggered cGAS activation through its Zn^2+^-binding activity. (A) HPLC analysis of 2′3′-cGAMP production in a reconstituted in vitro cGAS assay containing recombinant cGAS (10 μM), dsDNA (10^-2^ mg/mL), ATP (0.2 mM), and GTP (0.2 mM), with increasing amounts of recombinant ESAT-6 (0, 5, 10, 15, 20 μM) and Zn^2+^ (5 mM) add-back as indicated. (B) HPLC analysis of nucleotide products from the in vitro cGAS reaction performed in the presence of wild-type ESAT-6 (10 μM), heat-inactivated ESAT-6 (10 μM), or ESAT-6(H26A/D30A) (10 μM). Peaks corresponding to GTP, ATP, and 2′3′-cGAMP are indicated. (C) Confocal microscopy of cells stained with the Zn^2+^-sensitive probe Zinpyr-1 and transfected with Cy3-labeled ISD together with wild-type ESAT-6 or ESAT-6^H26A/D30A^, as indicated. Nuclei were stained with DAPI. (D) Confocal microscopy of HeLa cells expressing cGAS-eGFP and transfected with Cy3-labeled ISD in the presence or absence of wild-type ESAT-6 or ESAT-6(H26A/D30A). Nuclei were stained with DAPI. (E) Flow cytometric analysis of HeLa cells expressing cGAS-eGFP and/or Cy3-labeled ISD together with WT ESAT-6 or ESAT-6^H26A/D30A^, as indicated. (F) RT-qPCR analysis of gene expression in cells transfected with ISD and expressing wild-type ESAT-6 with Zn^2+^ supplementation as indicated. Gene expression was normalized to the indicated internal control and presented as relative expression values. *P* values were calculated using two-tailed Student’s t-test for paired comparisons or one-way or two-way ANOVA for multiple comparisons. Error bars represent ± SEM. \**P* < 0.05; \*\**P* < 0.01; \*\*\**P* < 0.001; ns, not significant.

Using the Zn^2+^-sensitive probe Zinpyr-1, we observed punctate intracellular Zn^2+^ signals following Cy3-ISD stimulation, whereas co-expression of wild-type ESAT-6 reduced this signal; the H26A/D30A mutant had a substantially weaker effect (Figure 5C). In HeLa cells expressing cGAS-eGFP, co-expression of wild-type ESAT-6 reduced Cy3-ISD-induced cGAS puncta formation, and this effect was attenuated in the H26A/D30A mutant (Figure 5D). Flow cytometry further showed that ESAT-6 had little effect on the intracellular Cy3-DNA signal itself (Figure 5E), supporting a mechanism distinct from CFP-10-mediated DNA depletion.

Functionally, wild-type ESAT-6 significantly attenuated ISD-induced expression of *Ifnb1* and many ISGs (Figures 5F and S6A). Consistent with a Zn^2+^-dependent mechanism, Zn^2+^ supplementation restored the induction of *Ifnb1* and ISGs in ESAT-6-expressing cells, and in a parallel assay, the ESAT-6^H26A/D30A^ mutant remained markedly less effective than WT ESAT-6 (Figure S6B). Together, these findings support a model in which ESAT-6 limits the Zn^2+^ conditions required for productive cGAS activation.

### ESAT-6 and CFP-10 act cooperatively to suppress cGAS–STING during infection

Having established that ESAT-6 and CFP-10 inhibit cGAS activation through distinct biochemical mechanisms, we next asked whether they act together during infection. Notably, analysis of WT *M. marinum* infection in *cGas^−/–^* and *Sting^−/–^* macrophages showed that the interferon program elicited by wild-type bacteria is itself largely dependent on cGAS-STING (Figure S7A). Consistent with the idea of biochemical cooperation, HPLC-based reconstitution assays showed that ESAT-6 and CFP-10 together suppressed dsDNA-triggered 2′3′-cGAMP production more efficiently than either protein alone, whereas combinations containing CFP-10^R57A^ or ESAT-6^H26A/D30A^ were markedly less effective (Figure S7B). Immunoblot analysis further showed that infection with the Δ*esat-6*Δ*cfp-10* strain induced stronger TBK1 phosphorylation than WT infection in C57 macrophages, whereas this difference was largely lost in *cGas^−/–^* and *Sting^−/–^*cells (Figure 6A).

**Figure 6.**
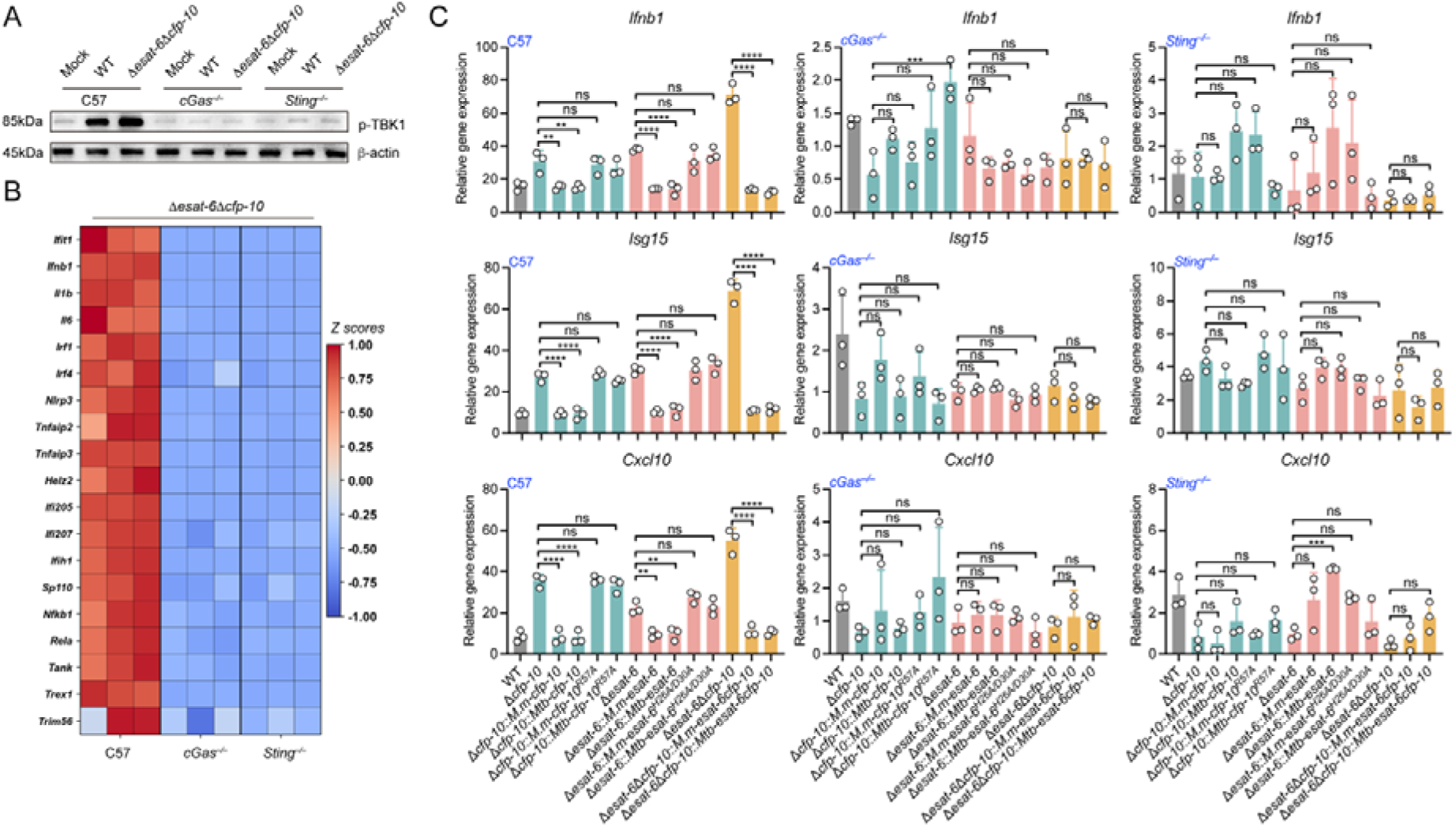
ESAT-6 and CFP-10 suppress innate immune activation through the cGAS-STING pathway during infection. (A) Immunoblot analysis of phosphorylated TBK1 (p-TBK1) in C57, *cGas^−/−^*, and *Sting^−/−^* macrophages infected with WT or Δ*esat-6*Δ*cfp-10* bacteria (MOI = 20, 4 h). β-actin was used as a loading control. Representative blot from 3 independent experiments. (B) Heatmap showing Z-score-normalized expression of selected immune-related genes from transcriptomic analysis of Δ*esat-6*Δ*cfp-10*-infected C57, *cGas^−/−^*, and *Sting^−/−^* macrophages (MOI = 20, 4 h). n = 3 independent biological replicates per group. (C) RT-qPCR analysis of gene expression in C57, *cGas^−/−^*, and *Sting^−/−^* macrophages infected with the indicated WT, deletion, and complementation strains, including WT or mutant *esat-6* and *cfp-10* alleles from *M. marinum* (*Mm*) or *M. tuberculosis* (*Mtb*) (MOI = 20, 4 h). Gene expression was normalized to the indicated internal control and presented as relative expression values. *P* values were calculated using two-tailed Student’s t-test for paired comparisons or one-way or two-way ANOVA for multiple comparisons. Error bars represent ± SEM. \**P* < 0.05; \*\**P* < 0.01; \*\*\**P* < 0.001; ns, not significant.

Pathway enrichment and differential expression analyses showed that transcripts preferentially induced by Δ*esat-6*Δ*cfp-10* infection in C57 macrophages, relative to *cGas^−/–^* and *Sting^−/–^* cells, were enriched for innate immune, cytokine, and cytosolic DNA-sensing pathways (Figure S8A–S8D). Consistently, heatmap analysis of representative response genes showed that the IFN- and inflammation-associated program induced by Δ*esat-6*Δ*cfp-10* infection in C57 macrophages was broadly attenuated in both knockout backgrounds (Figure 6B).

Genetic complementation further linked this phenotype to the biochemical activities of ESAT-6 and CFP-10. In WT C57 macrophages, complementation of Δ*cfp-10* with WT *cfp-10* from either *M. marinum* or *M. tuberculosis* restored suppression of *Ifnb1*, *Isg15*, and *Cxcl10*, whereas the DNA-processing-defective *cfp-10^R57A^* allele did not (Figure 6C). Similarly, complementation of Δ*esat-6* with WT *esat-6* from either species restored suppression, whereas the Zn^2+^-binding-defective *esat-6^H26A/D30A^* allele remained defective. The Δ*esat-6*Δ*cfp-10* double mutant induced the strongest response, which was reduced by *esat-6*/*cfp-10* co-complementation from *M. marinum* or *M. tuberculosis* (Figure 6C). Importantly, these strain-dependent differences were largely abolished in *cGas^−/–^*and *Sting^−/–^* macrophages, where deletion, WT complementation, and mechanism-defective complementation strains induced similarly low levels of *Ifnb1*, *Isg15*, and *Cxcl10* (Figure 6C).

This pattern was reproduced in RAW 264.7 macrophages (Figure S8E and S8F), and the enhanced induction of *Ifnb1* and ISGs by Δ*esat-6*Δ*cfp-10* strains was markedly reduced or absent in *cGas^−/–^*and *Sting^−/–^* macrophages (Figures S9A and S9B). Similar cGAS-STING dependence was observed in a *Corynebacterium pseudotuberculosis* model, which also encodes ESAT-6/CFP-10-like homologs (Figure S10A and S10B). Together, these data indicate that ESAT-6 and CFP-10 act cooperatively through conserved biochemical activities to suppress cGAS-STING-dependent innate immune activation during bacterial infection.

### ESAT-6 and CFP-10 promote virulence by suppressing cGAS-dependent immunity *in vivo*

To assess the in vivo relevance of ESAT-6/CFP-10-mediated immune suppression, we first used the zebrafish *M. marinum* infection model. Compared with WT infection, the Δ*esat-6*Δ*cfp-10* mutant markedly prolonged host survival and reduced bacterial burdens in immune organs by approximately two orders of magnitude, phenotypes comparable to those of the ESX-1-deficient Δ*mycP1* strain (Figures 7A and 7B). Despite their similar attenuation, only the Δ*esat-6*Δ*cfp-10* mutant elicited robust expression of *ifnphi1*, *isg15*, *tlr7*, *nod2*, *ripk2*, and *rela* (Figure 7C). Thus, the enhanced immune response to the double mutant cannot be explained solely by reduced bacterial burden or generalized ESX-1 dysfunction.

**Figure 7.**
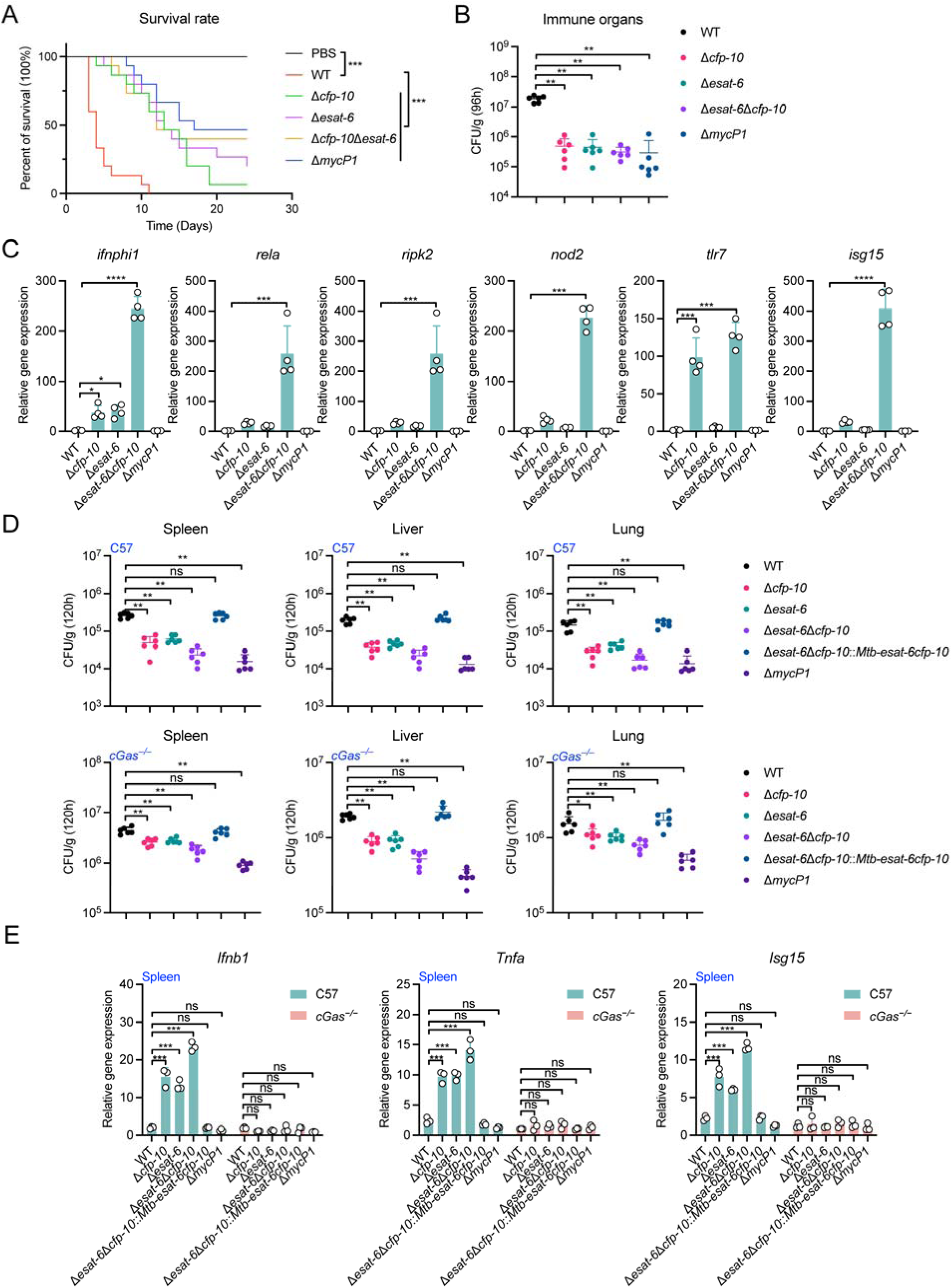
Loss of ESAT-6 and CFP-10 attenuates mycobacterial virulence while enhancing cGAS-dependent host immunity *in vivo*. (A) Kaplan-Meier survival analysis of zebrafish injected with PBS or infected with WT, Δ*esat-6,* Δ*cfp-10*, Δ*esat-6*Δ*cfp-10*, or Δ*mycP1* strains over the indicated time course (1 × 10^5^ CFU/fish). n = 15 fish per group. (B) Quantification of bacterial burdens in zebrafish immune organs at 96 h after infection, expressed as colony-forming units (CFU) per gram of tissue (1 × 10^5^ CFU/fish). n = 5 fish per group; each dot represents one fish. (C) RT-qPCR analysis of *ifnphi1*, *rela*, *isg15*, *tlr7*, *nod2*, and *ripk2* in zebrafish infected with the indicated strains (1 × 10^5^ CFU/fish). n = 4. (D) Quantification of bacterial burdens in spleen, liver, and lung from C57 and *cGas^−/−^* mice infected with the indicated strains, expressed as CFU per gram of tissue at 120 h after infection (1 × 10^5^ CFU/mouse). n = 5 mice per group; each dot represents one mouse. (E) RT-qPCR analysis of gene expression in spleens from C57 and *cGas^−/−^* mice infected with the indicated strains (1 × 10^5^ CFU/mouse). n = 4 mice per group. *P* values were calculated using two-tailed Student’s t-test for paired comparisons or one-way or two-way ANOVA for multiple comparisons. Error bars represent ± SEM. \**P* < 0.05; \*\**P* < 0.01; \*\*\**P* < 0.001; ns, not significant.

We next examined the contribution of cGAS in mice. At 120 h post-infection, the Δ*esat-6*Δ*cfp-10* mutant exhibited substantially reduced burdens in the spleen, liver, and lung of C57BL/6 mice, whereas expression of *M. tuberculosis esat-6–cfp-10* restored colonization toward WT levels (Figure 7D). By contrast, in *cGas^−/–^* mice, the attenuation of the double mutant was markedly diminished, with organ bacterial burdens no longer showing the same degree of reduction seen in WT animals. Consistent with these findings, the marked induction of splenic *Isg15*, *Ifnb1*, and *Tnfa* by the double mutant in C57BL/6 mice was abolished in *cGas*^−/–^ animals and reversed by complementation (Figure 7E). Collectively, these findings demonstrate that ESAT-6 and CFP-10 promote mycobacterial virulence *in vivo* while specifically suppressing cGAS-dependent innate immune activation.

### The Δ*esat-6*Δ*cfp-10* strain confers enhanced protection against secondary challenge

Because the Δ*esat-6*Δ*cfp-10* strain combined strong attenuation with heightened host immune activation, we asked whether it could provide protective immune priming *in vivo*. Using a zebrafish immunization-challenge model, animals were first treated with PBS, WT *M. marinum*, or the Δ*esat-6*Δ*cfp-10* strain, and subsequently challenged with WT *M. marinum* (Figure 8A). Prior exposure to the Δ*esat-6*Δ*cfp-10* strain resulted in improved survival following WT challenge compared with control groups (Figure 8B).

**Figure 8.**
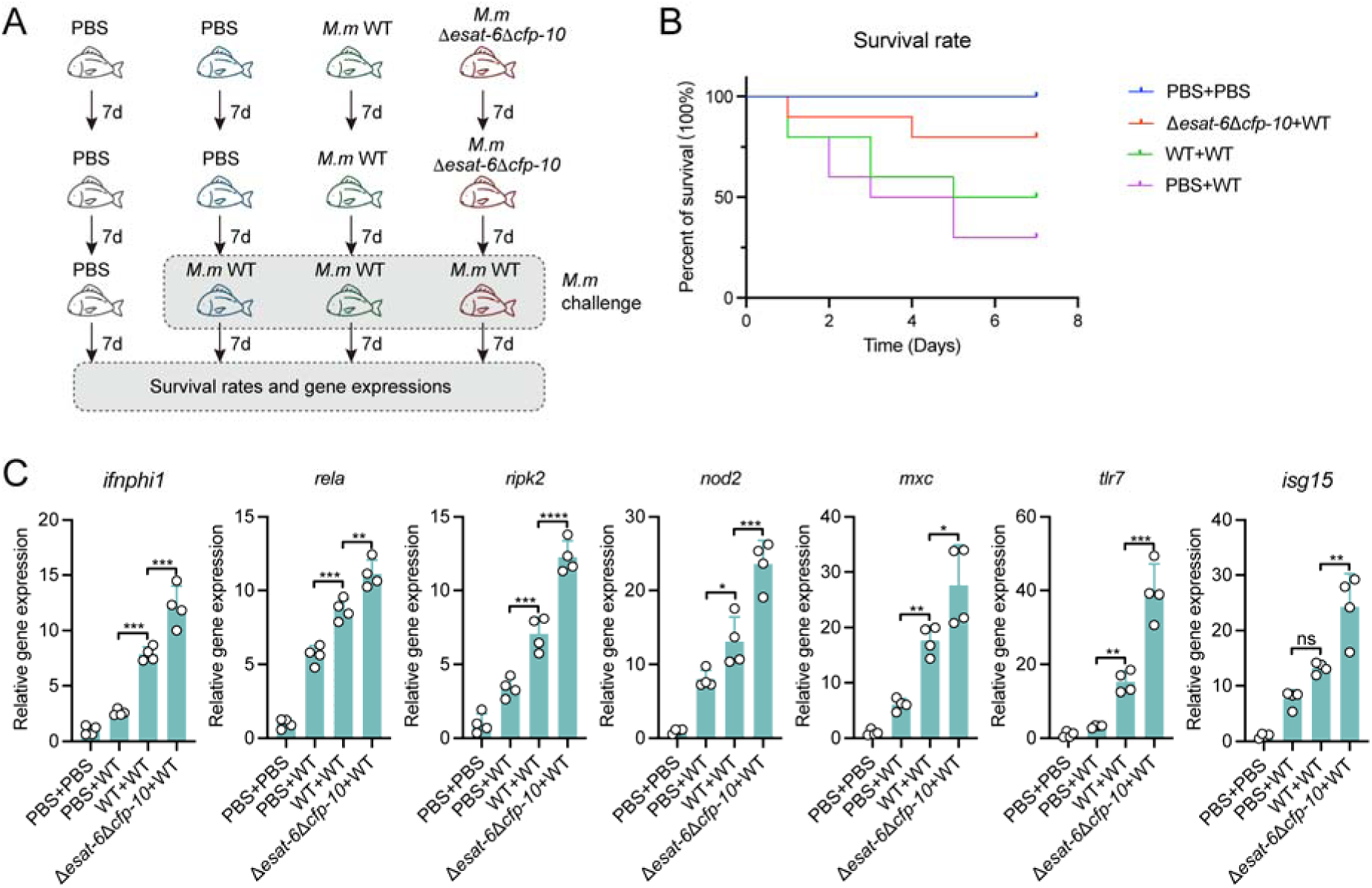
Protective priming induced by the Δ*esat-6*Δ*cfp-10* strain in a zebrafish challenge model. (A) Schematic of the zebrafish immunization-challenge experiment. Animals were treated sequentially with PBS, WT *M. marinum* (1 × 10^5^ CFU/fish), or the Δ*esat-6*Δ*cfp-10* strain (1 × 10^5^ CFU/fish) as indicated, followed by WT *M. marinum* challenge (5 × 10^5^ CFU/fish) and subsequent analysis of survival and gene expression. (B) Kaplan-Meier survival analysis of zebrafish subjected to the indicated immunization-challenge regimens. n = 10 fish per group. (C) RT-qPCR analysis of gene expression in zebrafish from the indicated experimental groups after WT challenge. n = 4 fish per group. *P* values were calculated using two-tailed Student’s t-test for paired comparisons or one-way or two-way ANOVA for multiple comparisons. Error bars represent ± SEM. \**P* < 0.05; \*\**P* < 0.01; \*\*\**P* < 0.001; ns, not significant.

Fish previously exposed to the double-mutant strain showed stronger induction of multiple immune-related genes after WT challenge than did PBS- or WT-pretreated controls (Figure 8C). This heightened secondary response is consistent with enhanced innate immune priming following prior exposure to the attenuated double-mutant strain. Together, these findings indicate that simultaneous disruption of ESAT-6 and CFP-10 not only reduces mycobacterial virulence *in vivo* but also preserves, and potentially enhances, the capacity to induce protective host responses upon subsequent challenge.

## DISCUSSION

Our findings redefine the ESX-1 system as one that not only grants mycobacterial DNA access to the cytosol but also deploys its major substrates to calibrate the intensity of the ensuing immune response. Extensive prior work has established that ESX-1, ESAT-6, and CFP-10 are required for phagosomal membrane damage, cytosolic access, secretion of additional ESX-1 substrates, and virulence^3,18,28,29^. That framework, however, left a central problem unresolved: if ESX-1-mediated membrane rupture exposes mycobacterial and host DNA to the cytosol, why does this not provoke overwhelming cGAS–STING activation^1,9,10,28,30^? Our findings do not replace this framework. Rather, they suggest that after secretion, ESAT-6 and CFP-10 act directly on components of the cytosolic environment that determine the magnitude of cGAS activation. In this expanded model, ESX-1-mediated membrane permeabilization promotes DNA exposure and pathway activation, whereas additional biochemical activities of individual substrates fine-tune the resulting response.

CFP-10 acts at the level of the activating ligand, reducing the availability of stimulatory DNA species required for productive cGAS activation. By functioning as a 5′–3′ exonuclease that preferentially processes linear DNA and requires R57 for this activity, CFP-10 appears to suppress host DNA sensing before the pathway is fully engaged. This ligand-centric strategy is particularly advantageous given the high local concentration of DNA that follows phagosomal rupture. ESAT-6, by contrast, operates through a fundamentally distinct mechanism: rather than depleting DNA, it restricts the Zn^2+^ conditions that support efficient cGAS activation^14,31,32^. Pathogen strategies that antagonize cGAS–STING commonly target DNA-sensing components, cyclic dinucleotide messengers, or downstream STING signaling^33–35^. Our data suggest that pathogens can also interfere with the local physicochemical environment required for innate immune receptor function. ESAT-6 therefore acts not on the ligand itself but on the ionic conditions that enable signal conversion.

This represents an unusual strategy in host–pathogen interactions: rather than competing for metals as nutrients, the pathogen sequesters a metal to disable a host sensor. Metal restriction has classically been framed as nutritional immunity, a host strategy that starves microbes of essential nutrients such as iron, zinc, and manganese^36^. Here, the logic is reversed: a pathogen-encoded effector locally restricts a metal ion not for its own metabolism but to disable a host immune sensor. This reframes trace metals as contested regulators of innate immune signal transduction, not merely nutritional resources. Whether other pathogens have evolved analogous strategies to manipulate metal-dependent host defenses can now be tested within this framework.

Together, the biochemical data support a bipartite model in which CFP-10 reduces ligand availability and ESAT-6 limits cofactor support for cGAS activation. These activities are mechanistically distinct and complementary in reconstituted and cell-based systems. A key conceptual implication is that efficient suppression of cGAS-dependent immunity cannot rely on DNA hydrolysis alone. Following ESX-1-mediated phagosomal breach, the cytosolic DNA load is substantial, and cGAS activation is driven by liquid–liquid phase separation, which concentrates ligand and sensor by orders of magnitude and can shield encapsulated DNA from cytosolic nucleases such as TREX1 ^5,14,37^. Moreover, host mitochondrial DNA released during infection can serve as an additional cGAS ligand. Thus, *M. tuberculosis* appears to deploy a layered evasion program: CFP-10 restricts the available DNA ligand, ESAT-6 constrains the Zn^2+^-dependent physicochemical conditions required for efficient cGAS activation, CdnP hydrolyzes the second messenger cGAMP, and MmsA targets STING for degradation ^34,38^. Our data place ESAT-6 and CFP-10 within this broader framework, showing that ESX-1 acts not only upstream of cytosolic DNA exposure but also at the level of signaling permissiveness.

The translational implications are immediate. BCG lacks RD1 and therefore lacks a functional ESX-1 system, a major basis for its attenuation but also a likely constraint on its ability to engage cytosolic immune pathways ^39^. Our data suggest an alternative design principle for attenuation: rather than abolishing cytosolic access entirely, one could selectively disable the immune-suppressive functions embedded within the ESX-1 program. Consistent with this idea, simultaneous disruption of ESAT-6 and CFP-10 strongly attenuated virulence while preserving robust immunostimulatory potential and enhanced protection in vivo. These findings support a model of attenuation by de-cloaking, in which removal of immune evasion, rather than wholesale loss of host engagement, yields a more favorable balance between safety and immunogenicity. More broadly, because the Zn^2+^-binding interface of ESAT-6 and the DNA-processing pocket of CFP-10 are biochemically discrete, they may represent tractable anti-virulence targets.

In summary, ESAT-6 and CFP-10 limit cGAS–STING signaling by reducing stimulatory DNA availability and restricting the Zn^2+^ conditions required for productive sensing, providing a mechanistic explanation for how virulent ESX-1-intact mycobacteria can open the phagosome yet keep innate activation in check. These findings expand the functional repertoire of ESAT-6 and CFP-10 alongside their established roles in secretion, phagosomal permeabilization, and cytosolic immune activation. More broadly, they indicate that the intensity of innate immune sensing is not fixed by the amount of ligand exposed, but is actively modulated at the interface between pathogen-encoded biochemistry and the physicochemical requirements of host sensors, a principle likely to extend beyond ESX-1 and cGAS.

## STAR METHODS

Detailed methods are provided in the online version of this paper and include the following:

- KEY RESOURCES TABLE
- RESOURCE AVAILABILITY

- Lead contact
- Materials availability
- Data and software availability
- EXPERIMENTAL MODEL AND SUBJECT DETAILS

- Study design
- Bacterial strains, cell lines, and culture conditions
- Plasmid construction and site-directed mutagenesis
- Generation of *M. marinum* mutants and complementation strains
- Preparation of competent cells and transformation
- Recombinant protein expression and purification
- Size-exclusion chromatography
- *In silico* analysis and structural modeling
- Electrophoretic mobility shift assay (EMSA)
- DNA-processing assays
- Metal dependence of CFP-10-mediated DNA processing
- Growth and toxicity assays
- Lentiviral packaging and transduction
- *In vitro* cGAS enzymatic reactions and HPLC analysis of 2′3′-cGAMP
- PAR assay for Zn^2+^ binding
- Confocal imaging of cGAS puncta and intracellular bioavailable Zn^2+^
- Flow cytometry
- Isolation of primary peritoneal macrophages
- Macrophage infection assays
- Intracellular bacterial load assay
- Immunoblotting
- RNA extraction, reverse transcription, and RT-qPCR
- Transcriptomic analysis
- Mouse infection experiments
- Zebrafish infection experiments
- Zebrafish immunization-challenge experiments
- QUANTIFICATION AND STATISTICAL ANALYSIS

- Statistics and reproducibility

## SUPPLEMENTAL INFORMATION

Supplemental information can be found online.

## ACKNOWLEDGEMENTS

This work was supported by the National Natural Science Foundation of China (32330004 to X.S.; 31970114 and 32170130 to Y.W.; 32670157 and 32270134 to L.X.), the Shaanxi Fundamental Science Research Project for Chemistry & Biology (22JHZ008 to X.S.), and the Natural Science Basic Research Program of Shaanxi (2026JC-YXQN-088 to L.X.), and the Ningbo Top Medical and Health Research Program (2023020713 to X.S.). We thank Zhengfan Jiang (Beijing University) for generously providing the *cGas^−/−^* and *Sting^−/−^* mice and HeLa cells. We also thank the Teaching and Research Core Facility at the College of Life Sciences (Ningjuan Fan, Xiyan Chen, and Hui Duan) and the Life Science Research Core Services, NWAFU (Luqi Li) for technical support.

## AUTHOR CONTRIBUTIONS

Conceptualization, L.X., Y.W., Y.S., and X.S.; Data curation, L.X., J.H., D.P., C.W., and T.C.; Formal analysis, L.X., J.H., D.P., C.W., T.C., and J.L.; Funding acquisition, L.X., Y.W., Y.S., and X.S.; Investigation, L.X., J.H., D.P., C.W., T.C., J.L., X.Z., S.L., and L.Z.; Methodology, L.X., J.H., D.P., Y.W., Y.S., and X.S.; Project administration, L.X., Y.W., Y.S., and X.S.; Resources, L.X., Y.W., Y.S., and X.S.; Supervision, L.X., Y.W., Y.S., and X.S.; Validation, L.X., J.H., D.P., C.W., T.C., and J.L.; Visualization, L.X., J.H., D.P., C.W., and T.C.; Writing – original draft, L.X., J.H.; Writing – review and editing, L.X., J.H., D.P., Y.W., Y.S., and X.S. All authors discussed the results and commented on the manuscript.

## COMPETING INTERESTS

The authors declare no competing interests.

## KEY RESOURCES TABLE

| REAGENT or RESOURCE | SOURCE | IDENTIFIER |
| --- | --- | --- |
| **Chemicals, peptides, and recombinant proteins** |  |  |
| NaCl | Sigma-Aldrich | CAS: 7647-14-5 |
| MgCl<sub>2</sub> | Sigma-Aldrich | CAS: 7786-30-3 |
| CaCl<sub>2</sub> | Sigma-Aldrich | CAS: 10043-52-4 |
| ZnCl<sub>2</sub> | Sigma-Aldrich | CAS: 7646-85-7 |
| FeCl<sub>3</sub> | Sigma-Aldrich | CAS: 7705-08-0 |
| HCl | Sigma-Aldrich | CAS: 7647-01-0 |
| Agarose | Sigma-Aldrich | CAS: 9012-36-6 |
| 2-Mercaptoethanol | Sigma-Aldrich | CAS: 60-24-2 |
| Adenosine 5'-triphosphate (ATP) disodium salt hydrate | Sigma-Aldrich | CAS: 34369-07-8 |
| Guanosine 5'-triphosphate (GTP) sodium salt hydrate | Sigma-Aldrich | CAS: 56001-37-7 |
| 2'3'-cGAMP sodium salt | Sigma-Aldrich | CAS: 1441190-66-4 |
| Sodium pyruvate solution | Sigma-Aldrich | CAS: 113-24-6 |
| HEPES solution | Sigma-Aldrich | CAS: 7365-45-9 |
| DNase I | Sigma-Aldrich | Cat. no.: 10104159001 |
| Tris-base | Sigma-Aldrich | CAS: 77-86-1 |
| DTT | Sigma-Aldrich | CAS: 3483-12-3 |
| Acrylamide | Sigma-Aldrich | CAS: 79-06-1 |
| EDTA | Sigma-Aldrich | CAS: 60-00-4 |
| BSA | Sigma-Aldrich | CAS: 9048-46-8 |
| SDS | Sigma-Aldrich | CAS: 151-21-3 |
| APS | Sigma-Aldrich | CAS: 7727-54-0 |
| TEMED | Sigma-Aldrich | CAS: 7677-21-6 |
| IPTG | Sigma-Aldrich | CAS: 367-93-1 |
| Tween 80 | Sigma-Aldrich | CAS: 9005-65-6 |
| Kanamycin | Sigma-Aldrich | CAS: 25389-94-0 |
| Ampicillin | Sigma-Aldrich | CAS: 69-53-4 |
| L-Glutamine (100×) | Beyotime | CAS: 56-85-9 |
| 0.25% Trypsin | Beyotime | Cat. no.: C0201 |
| 0.05% Trypsin | Beyotime | Cat. no.: C0202 |
| DAPI | Beyotime | Cat. no.: C1002 |
| λDNA | Thermo Scientific | Cat. no.: SD0021 |
| Transfection reagent | Yeasen | Cat. no.: 40808ES02 |
| Zinpyr-1 | Yeasen | Cat. no.: 735224ES03 |
| 4-(2-Pyridylazo)resorcinol (PAR) | Sigma-Aldrich | CAS: 1141-59-9 |
| **Media and solutions** |  |  |
| Phosphate-Buffered Saline (PBS) | Beyotime | #C0221A |
| Penicillin-Streptomycin Solution (100×) | Beyotime | #C0222 |
| Dulbecco's Modified Eagle Medium (DMEM), high glucose | GIBCO | #11965-084 |
| Opti-MEM I Reduced Serum Medium | GIBCO | #11058-021 |
| RPMI 1640 | GIBCO | #11875-093 |
| Fetal bovine serum (FBS) | Beyotime | #C0234 |
| Sodium pyruvate Solution (100 mM) | Sigma-Aldrich | #S8636; CAS: 113-24-6 |
| HEPES Solution (1 M) | Sigma-Aldrich | #H0887; CAS: 7365-45-9 |
| MEM Non-Essential Amino Acids (100×) | GIBCO | #11140-050 |
| 2-Mercaptoethanol | Sigma-Aldrich | #M6250; CAS: 60-24-2 |
| QuickBlock™ Blocking Buffer | Beyotime | #P0252 |
| QuickBlock™ Primary Antibody Dilution Buffer for Western Blot | Beyotime | #P0256 |
| RIPA Lysis Buffer | Beyotime | #P0013B |
| **Bacterial strains** |  |  |
| *M. marinum* | Our laboratory | N/A |
| *E. coli* DH5α | Our laboratory | N/A |
| *Corynebacterium pseudotuberculosis* | Our laboratory | N/A |
| *M. marinum* mutant strains (see Table S1) | Our laboratory | N/A |
| **Antibodies** |  |  |
| Mouse monoclonal anti-β-actin (clone C4) | Santa Cruz Biotechnology | #sc-47778, RRID: AB_2714189 |
| Rabbit monoclonal anti-Phospho-TBK1/NAK (Ser172) (clone D52C2) | Cell Signaling Technology | #5483, RRID: AB_10693472 |
| **Experimental Models: Organisms/Strains** |  |  |
| Mouse: C57BL/6 | SPF | N/A |
| Mouse: *cGas*<sup>-/-</sup> | Dr. Zhengfan Jiang | N/A |
| Mouse: *Sting*<sup>-/-</sup> | Dr. Zhengfan Jiang | N/A |
| **Experimental Models: Cell lines** |  |  |
| Mouse: RAW 264.7 cells | ATCC | #TIB-71 |
| Human: THP-1 | Beyotime | #C6960 |
| Human: HeLa | N/A | 40 |
| **Commercial assays** |  |  |
| RNAeasy™ Animal RNA Isolation Kit with Spin Column | Beyotime | #R0027 |
| RNAprep Pure Tissue Kit | TIANGEN | #DP431 |
| RNAprep Pure Cell/Bacteria Kit | TIANGEN | #DP430 |
| NucleoBond® Xtra Midi Plus Kit | Macherey-Nagel | #740422.50 |
| FastKing RT Kit (With gDNase) | TIANGEN | #KR116-02 |
| RealUniversal Color PreMix (SYBR Green) | TIANGEN | #FP201-02 |
| Primers for qRT-PCR (see Table S3) | This paper | N/A |
| **Other** |  |  |
| GraphPad Prism 8.01 | GraphPad | https://www.graphpad.com/ |
| **Deposited data** |  |  |
| RNA-seq | CNSA (China National GeneBank Sequence Archive) | Accession number: CNP0009381 |

## RESOURCE AVAILABILITY

### Lead contact

Further information and requests for resources and reagents should be directed to and will be fulfilled by the lead contact, Xihui Shen.

### Materials availability

This study did not generate new, unique reagents.

### Data and software availability

The data supporting the findings of this study (including RNA-seq FASTQ files) have been deposited into CNSA (China National GeneBank Sequence Archive) with accession number CNP0009381 (http://db.cngb.org/cnsa/project/CNP0009381_c9f5430e/reviewlink/).

## EXPERIMENTAL MODEL AND METHOD DETAILS

### MATERIALS AND METHODS

#### Study design

This study was designed to define how the ESX-1 effectors CFP-10 and ESAT-6 regulate host cGAS-STING signaling during mycobacterial infection. We combined biochemical reconstitution assays, genetic analysis in *Mycobacterium marinum*, cell-based infection models, fluorescence imaging, flow cytometry, and animal infection and immunization models to determine how CFP-10-mediated DNA processing and ESAT-6-mediated Zn^2+^ sequestration converge on host DNA sensing. Because long-term BSL-3 experiments with *Mycobacterium tuberculosis* (*Mtb*) were not available in our setting, *M. marinum* was used as a BSL-2 surrogate model, while selected *Mtb* homologs were included in complementation and biochemical experiments where indicated. Animal experiments were approved by the Animal Welfare and Research Ethics Committee of Northwest A&F University (protocol XN2023-1004).

#### Bacterial strains, cell lines, and culture conditions

The bacterial strains, plasmids, cell lines, and selected primers used in this study are listed in Tables S1 and S2. *E. coli* DH5α was used for cloning and BL21 (DE3) for recombinant protein expression. *M. marinum* wild-type (WT), single-gene deletion mutants, including Δ*esat-6,* Δ*cfp-10,* Δ*mycP1,* and Δ*eccCa_1_,* the Δ*esat-6*Δ*cfp-10* double mutant, and their corresponding complementation strains were used for infection experiments. For *Corynebacterium pseudotuberculosis* (*C. pseudotuberculosis*, CP), the WT strain and the Δ*esat-6*Δ*cfp-10* double mutant were used for infection experiments. *E. coli* strains were cultured in Luria-Bertani (LB) medium (Sangon Biotech, A507002, China) at 37°C, and *M. marinum* strains were cultured at 30°C in Middlebrook 7H9 broth (BD Difco, 271310, USA) supplemented with ADC (Hopebiol, HB9233a, China), glycerol, and Tween-80 (Solarbio, T8360, China), or on 7H10 agar (BD Difco, 262710, USA) supplemented with OADC (Hopebiol, HB6271a, China) and glycerol. *C. pseudotuberculosis* ATCC 19410 was cultured at 37°C in brain heart infusion (BHI) broth (Solarbio, LA0360, China). Kanamycin and ampicillin were used at 25 μg/mL and 50 μg/mL, respectively, when required. Nalidixic acid was used at a final concentration of 20 μg/mL when required. RAW 264.7 macrophages (ATCC, #TIB-71), HEK293T cells (ATCC, CRL-1126), and HeLa cells (ATCC, CRL-2) were maintained in high-glucose DMEM (Mishu, MI00622, China) supplemented with 10% fetal bovine serum (GIBCO, #10099-141, USA) and 1% penicillin-streptomycin (Beyotime, C0222, China). Primary peritoneal macrophages were maintained in RPMI 1640 (Mishu, MI00620, China) supplemented with 10% fetal bovine serum, 1% penicillin-streptomycin, nonessential amino acids (GIBCO, #11140-050, USA), sodium pyruvate (Sigma-Aldrich, 113-24-6, USA), HEPES (Sigma-Aldrich, 7365-45-9, USA), and β-mercaptoethanol (Sigma-Aldrich, 60-24-2, USA).

#### Plasmid construction and site-directed mutagenesis

Coding sequences corresponding to *esat-6*/*cfp-10* or their *Mtb* homologs (*rv3875*/*rv3874*) were amplified from *M. marinum* genomic DNA or synthesized DNA templates and inserted into shuttle or expression vectors using standard restriction enzyme-based cloning. pMV261-derived constructs were used for complementation in *M. marinum*. For *C. pseudotuberculosis*, the upstream and downstream homologous arms were fused by PCR using the *cfp-10*-up-F/*esat-6*-down-R primer pair and subsequently cloned into the pK18mobsacB plasmid to generate the *esat-6/cfp-10* deletion suicide plasmid pK18mobsacBΔ*esat-6*Δ*cfp-10*. pGEX6P-1-derived constructs were used to express GST-tagged CFP-10/Rv3874/ESAT-6/Rv3875 and their mutants, whereas pET28a-derived constructs were used to express His-tagged CFP-10/Rv3874/ESAT-6/Rv3875 and their mutant derivatives. Lentiviral constructs were generated in pLenti-CMV vectors for ectopic expression in mammalian cells. For fluorescence imaging, cGAS-eGFP, ESAT-6 and CFP-10-related expression plasmids were generated in pCMV-based mammalian vectors. Point mutations were introduced by overlap PCR and verified by sequencing. The principal mutants used in this study were CFP-10/Rv3874 R57A, which disrupts DNA interaction and DNA processing, and ESAT-6/Rv3875 H26A/D30A, which disrupts Zn^2+^ binding.

#### Generation of *M. marinum* mutants, *Corynebacterium pseudotuberculosis* mutants, and complementation strains

Single-gene deletion strains lacking *esat-6* or *cfp-10*, as well as the Δ*esat-6*Δ*cfp-10* double mutant, were generated in *M. marinum* using a modified CRISPR-Cas12a-based strategy. All *M. marinum* mutant strains were constructed by the Institute of Pathogen Biology, Chinese Academy of Medical Sciences, and preserved by the Agricultural and Environmental Microbiology Team at Northwest A&F University. Complementation strains were generated by introducing wild-type or mutant alleles into the corresponding deletion background using pMV261-derived plasmids, followed by selection on kanamycin-containing 7H10 plates. Strain identities were verified by PCR and/or sequencing-based validation. For *C. pseudotuberculosis* ATCC 19410, pK18mobsacBΔ*esat-6*Δ*cfp-10* was electroporated into competent ATCC 19410 cells and plated on BHI agar containing kanamycin (25 μg/mL). Single-crossover strains were identified using the *cfp-10*-up-F/*esat-6*-down-R primer pair, followed by serial passage on BHI agar containing 20% sucrose. Double-crossover deletion mutants showing only the 1,319-bp PCR fragment were screened and designated ATCC 19410Δ*esat-6*Δ*cfp-10*.

#### Preparation of competent cells and transformation

Chemically competent *E. coli* DH5α and BL21 (DE3) cells were prepared using a CaCl_2_-based method. Briefly, log-phase cells were harvested, washed, incubated in cold 0.1 M CaCl_2_, resuspended in 0.1 M CaCl_2_ containing 15% glycerol, and stored at −80°C. Plasmids were introduced into *E. coli* by heat shock. Electrocompetent *M. marinum* cells were prepared by repeated washes in 10% glycerol/Tween-80-containing buffer and transformed by electroporation at 2.5 kV, followed by recovery in 7H9 broth and plating on selective 7H10 agar. For *C. pseudotuberculosis,* competent cells were washed four times with 10% glycerol and transformed by electroporation at 2.5 kV, followed by the addition of BHI medium, heat shock at 46°C for 6 min, and static recovery at 37°C. Standard plasmid miniprep, genomic DNA extraction, gel extraction, restriction digestion, and ligation procedures were performed as described ^41,42^.

#### Recombinant protein expression and purification

Expression plasmids were transformed into *E. coli* BL21 (DE3), and single colonies were inoculated into LB medium containing the appropriate antibiotic. Cultures were grown at 37°C to OD_600_ ∼0.3 and induced with 0.5 mM IPTG, then shifted to low temperature and incubated at 4°C with shaking for 12–14 h to promote soluble expression. Bacterial pellets were resuspended in ice-cold binding buffer and lysed by high-pressure disruption. Soluble lysates were clarified by centrifugation and applied to the affinity resin. GST-tagged proteins were purified using glutathione affinity chromatography with a binding/wash buffer containing 137 mM NaCl and 50 mM Tris-HCl (pH 7.4), followed by elution in the same buffer supplemented with reduced glutathione. His-tagged proteins were purified using Ni-NTA chromatography with phosphate-based buffers containing imidazole ^41,42^. Eluted proteins were dialyzed into storage buffer containing salt, buffer, and glycerol, quantified spectrophotometrically, and analyzed by SDS-PAGE with Coomassie staining before use in biochemical assays.

#### Size-exclusion chromatography

To further improve protein purity and homogeneity, affinity-purified ESAT-6 and CFP-10 proteins were subjected to size-exclusion chromatography. Briefly, eluted proteins from Ni-NTA or glutathione affinity purification were first concentrated using centrifugal ultrafiltration devices and then loaded onto a pre-equilibrated Superdex 75 Increase 10/300 GL column (GE Healthcare, Uppsala, Sweden) connected to an ÄKTA pure chromatography system (GE Healthcare, Uppsala, Sweden). The column was equilibrated with gel filtration buffer containing 20 mM Tris-HCl (pH 7.4), 150 mM NaCl, 1 mM DTT, and 5% glycerol. Protein samples were centrifuged at high speed and passed through a 0.22-μm filter before loading to remove aggregates and insoluble material. Chromatographic separation was performed at 4°C with a flow rate of 0.5 mL/min, and protein elution was monitored by absorbance at 280 nm. Fractions corresponding to the major monodisperse peak were collected and analyzed by SDS-PAGE. Fractions containing highly purified ESAT-6 or CFP-10 were pooled, concentrated as needed, flash-frozen in liquid nitrogen, and stored at −80°C until use in downstream biochemical and cell-based assays.

#### *In silico* analysis and structural modeling

Protein sequence conservation analysis was performed using ESAT-6/CFP-10 homologs from related mycobacterial species. Putative DNA-binding or Zn^2+^-binding surfaces were inferred from structural modeling (https://alphafold.com/) and molecular docking analyses (https://aideepmed.com/). Candidate functional residues were prioritized on the basis of sequence conservation and predicted spatial positioning relative to the modeled ligand-interaction surface. These analyses guided the design of the CFP-10^R57A^ and ESAT-6^H26A/D30A^ mutants used throughout the study.

#### Electrophoretic mobility shift assay (EMSA)

EMSA was performed to assess CFP-10/Rv3874-DNA interaction. Recombinant CFP-10 protein at the indicated concentrations was incubated with 40 ng of 100-bp double-stranded DNA in reaction buffer containing 20 mM Tris, 100 mM NaCl, 4 mM MgCl_2_, 1 mM DTT, and 10% glycerol (pH 7.4) at 37°C for 30 min. Reactions were mixed with loading buffer containing 0.5% SDS and resolved on 6% native polyacrylamide gels in 0.5 × TBE. Gels were stained with SYBR Safe, washed, and imaged. In these experiments, CFP-10 did not generate a stable retarded protein-DNA complex, but instead produced a progressive loss of the free DNA signal, which prompted further examination of DNA-processing activity.

#### DNA-processing assays

DNA-processing activity was assessed by incubating recombinant CFP-10/Rv3874 proteins with linear or circular DNA substrates in reaction buffer containing 20 mM Tris, 100 mM NaCl, 4 mM MgCl_2_, 1 mM DTT (Sigma-Aldrich, 3483-12-3, USA), 10% glycerol, and 1% NP-40 (pH 7.4) (Beyotime, P0013F, China) at 37°C for 30–60 min. GST protein and BSA were used as negative controls, and DNase I served as a positive control. Reaction products were resolved on 1% agarose gels and visualized using nucleic acid dye. To assess terminal preference, linear double-stranded DNA substrates bearing 5′ overhangs, 3′ overhangs, or protruding termini at both ends were compared. These experiments were used to define the substrate preference and 5′–3′ exonuclease-like behavior of CFP-10.

#### Metal dependence of CFP-10-mediated DNA processing

To test ion dependence, standard DNA-processing reactions were performed with 20 ng of λDNA (Thermo Scientific, SD0021, China) and 10 μM recombinant Rv3874 or CFP-10 protein in the presence or absence of EDTA (25 mM), followed by the addition of individual metal ions as indicated. The final ion concentrations were 4 mM Na^+^, 4 mM Mg^2+^, 1 mM Fe^3+^, 4 mM Ca^2+^, and 1 mM Zn^2+^. After incubation under standard reaction conditions, DNA degradation was analyzed by agarose gel electrophoresis.

#### Growth and toxicity assays

To evaluate possible toxicity associated with CFP-10 expression, growth assays were performed in both *E. coli* and *M. marinum*. In *E. coli*, BL21 (DE3) cells harboring pET28a-*cfp-10*, pET28a-*cfp-10^R57A^*, or empty vector were grown to OD_600_ ∼0.3 and induced with 0.5 mM IPTG, after which growth was monitored hourly by OD_600_ measurement. Growth curves of other relevant bacterial strains were determined as follows: bacterial strains were inoculated into the corresponding liquid medium at a 1:100 dilution to achieve an initial OD_600_ below 0.1. *E. coli* was measured every 1–2 h, while *M. marinum* was measured every 4–24 h. Measurements were continued until the cultures reached the stationary phase.

#### Lentiviral packaging and transduction

For ectopic expression of CFP-10/Rv3874, ESAT-6/Rv3875, or their mutants in mammalian cells, lentiviral particles were produced in HEK293T cells using a three-plasmid system consisting of transfer plasmid, psPAX2, and pMD2.G at a mass ratio of 4:3:1. HEK293T cells were seeded in 10-cm dishes and transfected with plasmid-transfection reagent (Yeasen, 40808ES02, China) complexes prepared in serum-free DMEM. Viral supernatants were collected at 48 h and 72 h after transfection, filtered through 0.45-μm membranes, and concentrated by ultracentrifugation (25,000 rpm, 2 h) (CP100WX, Hitachi Ltd., Japan) at 4°C. RAW 264.7 cells were transduced with concentrated virus and collected for downstream RNA expression analysis.

#### *In vitro* cGAS enzymatic reactions and HPLC analysis of 2′3′-cGAMP

cGAS activity was evaluated in a biochemical reconstitution system followed by HPLC (Alliance HPLC, Waters, USA) detection of 2′3′-cGAMP (Sigma-Aldrich, 1441190-66-4, USA) ^43^. The standard reaction buffer contained 0.2 mM ATP (Sigma-Aldrich, 34369-07-8, USA), 0.2 mM GTP (Sigma-Aldrich, 56001-37-7, USA), 100 mM NaCl, 40 mM Tris-HCl (pH 7.5), 1 mM MnCl_2_, 5 mM MgCl_2_, and 10^−2^ mg/mL double-stranded DNA (dsDNA). For assays examining Zn^2+^-facilitated cGAS activation, 5 mM ZnCl_2_ was additionally included. All reaction solutions were passed through a 0.22-μm filter before use. Purified cGAS was added at a final concentration of 10 μM, and recombinant CFP-10, ESAT-6, or the corresponding mutant proteins were added as indicated. Reactions were incubated at 37°C for 2 h.

Reactions were terminated by heating at 100°C for 20 min, followed by centrifugation at 12,000 rpm for 10 min to pellet denatured proteins. The resulting supernatants were transferred to HPLC sample vials and analyzed using an Alliance HPLC system. The mobile phase consisted of 150 mM Na_2_HPO_4_ (pH 5.2). This assay was used to quantify the effect of CFP-10 on DNA-triggered cGAS activation, the effect of ESAT-6 on Zn^2+^-facilitated cGAS activation, and the cooperative inhibition produced by ESAT-6 and CFP-10 together.

#### PAR assay for Zn^2+^ binding

Zn^2+^ binding by ESAT-6 was assessed using a PAR-based assay ^44^. The detection solution contained 50 mM Tris-HCl, 10 μM PAR (Sigma-Aldrich, 1141-59-9, USA), and 5 μM Zn^2+^ (pH 7.4). Recombinant ESAT-6 or ESAT-6^H26A/D30A^ was added at final concentrations of 0.4 to 2.5 μM, and samples without protein served as controls. After incubation at room temperature for 30 min, 200 μL of each reaction was transferred to a transparent 96-well plate, and absorbance spectra were recorded from 350 to 600 nm using a microplate reader. Free PAR and the (PAR)-Zn^2+^ complex show maximal absorbance at 410 nm and 500 nm, respectively, and changes in the absorbance profile were used to assess Zn^2+^ binding. Each assay included at least three technical replicates and was repeated independently at least three times.

#### Isothermal titration calorimetry (ITC)

Metal binding by ESAT-6 was further quantified by ITC (TA-Waters LLC, USA), essentially as described ^45^. The DNA-binding ability of CFP-10 was also assessed using synthetic 100-bp double-stranded DNA as the substrate. The dialysis buffer contained no EDTA or additional divalent cations such as Mg^2+^. Wild-type ESAT-6 and the H26A/D30A mutant were diluted to 25 μM in dialysis buffer, and 1 mM ZnCl_2_, CaCl_2_, CuCl_2_, or FeCl_3_ prepared in the same buffer was used as titrant. Before measurement, protein samples, ion solutions, and buffer were thoroughly mixed and degassed. CFP-10 was prepared under the same conditions, with cGAS included as a control; both proteins were diluted to 25 μM, and synthetic 100-bp dsDNA was used at a concentration of 500 μM. ITC experiments were performed by loading 1 mL of protein solution into the sample cell and 1 mL of dialysis buffer into the reference cell, followed by 25 sequential injections of 10 μL each, for a total titration volume of 250 μL. Control titrations were performed by injecting ion solutions into buffer alone under identical conditions, and dilution heats were subtracted during analysis. Data were analyzed using NanoAnalyze software to determine the binding constant (*Kd*), stoichiometry (n), and thermodynamic parameters (Δ*H*, Δ*G*, and Δ*S*). All experiments were repeated at least three times independently.

#### Confocal imaging of cGAS puncta and intracellular bioavailable Zn^2+^

HeLa cells were dissociated with 0.25% trypsin (Beyotime, C0201, China), counted, and seeded into glass-bottom confocal dishes at 30–40% confluence. For cGAS imaging, cells were cotransfected with pCMV-cGAS-eGFP and the indicated plasmids. After 20 h, the medium was replaced and cells were transfected with Cy3-labeled ISD for 4 h. Before imaging, cells were washed three times with PBS, stained with DAPI (Beyotime, C1002, China) for 5–10 min at room temperature in the dark, washed again, and maintained in PBS. Confocal microscopy (TCS SP8 SR, Leica Microsystems, Germany) was performed using 488 nm excitation for eGFP, 562 nm for Cy3, and 405 nm for DAPI.

For Zn^2+^ imaging, HeLa cells were seeded into glass-bottom dishes and transfected with the indicated plasmids, followed by Cy3-labeled ISD transfection 4 h before imaging. Cells were then stained with 2 μM Zinpyr-1 (Yeasen, 735224ES03, China) in PBS at 37°C for 20–30 min in the dark, washed, counterstained with DAPI, and imaged by confocal microscopy using 488 nm for Zinpyr-1, 562 nm for Cy3, and 405 nm for DAPI. Images were analyzed using ImageJ or microscope-associated software to quantify puncta formation, fluorescence intensity, and colocalization.

#### Flow cytometry

Flow cytometry was used to quantify intracellular fluorescent DNA and cGAS-associated fluorescence signals. Cells subjected to the imaging workflow were detached with trypsin, pelleted, resuspended in PBS, and filtered through nylon mesh before acquisition on a CytoFLEX flow cytometer (Beckman Coulter, China). Forward and side scatter were used to gate viable cells, after which fluorescence thresholds were defined using negative and positive controls. Typically, 1 × 10 cells were collected per sample. Data were analyzed using CytExpert (2.3.0.84).

#### Isolation of primary peritoneal macrophages

Primary peritoneal macrophages were isolated from C57BL/6, *cGas^−/−^*, and *Sting^−/−^* mice. To enrich peritoneal macrophages, mice were injected intraperitoneally with starch broth once daily for 3 consecutive days. One day after the final injection, mice were euthanized and the peritoneal cavity was lavaged repeatedly with sterile PBS. Recovered cells were pelleted, resuspended in RPMI complete medium, and plated in 24-well plates. After adherence, nonadherent cells were removed by medium exchange, and macrophages were used for infection experiments.

#### Macrophage infection assays

For infection experiments, RAW 264.7 cells or primary peritoneal macrophages were seeded in antibiotic-free complete medium such that cells reached ∼80% confluence on the day of infection. Mid-log *M. marinum* cultures were washed and resuspended in antibiotic-free cell culture medium. Cells were infected at a multiplicity of infection (MOI) of 20. Mid-log-phase *C. pseudotuberculosis* cultures were similarly washed and resuspended in antibiotic-free cell culture medium, and cells were infected at an MOI of 10. To synchronize infection, plates were centrifuged briefly at low speed and then incubated for 2 h to allow bacterial entry. This time point was defined as 0 h post-infection. Cells were washed with PBS to remove extracellular bacteria, replenished with antibiotic-containing complete medium, and incubated for an additional 4 h before harvest. RNA was extracted for RT-qPCR or transcriptomic analysis, and protein lysates were collected for immunoblotting where indicated. Wild-type, single-mutant, double-mutant, and complementation strains were compared side-by-side, including experiments in *cGas^−/−^*and *Sting^−/−^* macrophages.

#### Intracellular bacterial load assay

To determine the intracellular survival or proliferation of different bacterial strains, RAW 264.7 cells were infected with the indicated strains. To synchronize infection, the culture dishes were briefly centrifuged at low speed and then incubated for 2 h to allow bacterial invasion of the cells. This time point was defined as 0 h post-infection. The cells were then washed with PBS to remove extracellular bacteria, replenished with complete medium, and further incubated. Samples were collected at 0, 1, and 2 h. Host cells were lysed with an appropriate volume of 0.1% Triton X-100 to release intracellular bacteria. The lysates were serially diluted and plated onto the appropriate solid medium. After incubation until single colonies formed, the colonies were counted. Intracellular bacterial loads were expressed as colony-forming units per well, namely CFU/well. Biological replicates were included for each strain, and CFU values at different time points were statistically analyzed.

#### Immunoblotting

Primary peritoneal macrophages (PMs) from wild-type, *cGas^−/−^*, and *Sting^−/−^* mice were seeded in 6-well plates and infected with *M. marinum* (WT, Δ*esat-6*, Δ*cfp-10*, Δ*esat-6*Δ*cfp-10*, and Δ*mycP_1_*) at a multiplicity of infection (MOI) of 20. To synchronize infection, plates were centrifuged at 300 × g for 5 min. At the indicated time points, cells were washed with cold PBS and lysed in RIPA buffer (Beyotime, P0013, China) supplemented with protease and phosphatase inhibitors (Beyotime, P1045, China). Clarified lysates were quantified by BCA assay, mixed with 5× SDS loading buffer, and separated on 10% SDS-PAGE gels, followed by transfer to PVDF membranes (Millipore, FFP39, USA). Membranes were blocked in 5% BSA/TBST and incubated with antibodies against phospho-TBK1 (Ser172) (Cell Signaling Technology, #5483, USA) and β-actin (Santa Cruz Biotechnology, #sc-47778, USA), followed by HRP-conjugated secondary antibodies. Signals were visualized using ECL (Beyotime, P0018S, China) and quantified with ImageJ.

#### RNA extraction, reverse transcription, and RT-qPCR

Total RNA was extracted from cultured cells using the Beyotime animal RNA isolation kit (R0027) according to the manufacturer’s instructions. First-strand cDNA was synthesized using a reverse transcription kit (TransGen Biotech, AE311-02, China) according to the manufacturer’s instructions. Quantitative PCR was performed using SYBR-based chemistry (TransGen Biotech, AQ601-01-V2, China) on a real-time PCR system under the following conditions: 95°C for 30 s; 40 cycles of 95°C for 15 s and 58°C for 15 s; followed by melting curve analysis. Relative gene expression was normalized to *Actb*.

#### Transcriptomic analysis

For transcriptomic profiling, primary peritoneal macrophages from wild-type, *cGas^−/−^*, or *Sting^−/−^* mice were infected with *M. marinum* WT or Δ*esat-6*Δ*cfp-10* at an MOI of 20 for 4 h, with uninfected cells as controls. Total RNA was extracted using TRIzol (Invitrogen, 15596018CN, China), and samples with RIN ≥ 7.0 were used for poly(A)-enriched library preparation and Illumina PE150 sequencing. Raw reads were quality-checked and trimmed, aligned to the mouse reference genome using HISAT2, and quantified at the gene level using HTSeq. Differential expression analysis was performed using DESeq2 or edgeR, with |log2FC| ≥ 1 and FDR < 0.1 used as cutoffs. GO, KEGG, and GSEA were used to identify enriched pathways, with particular focus on cytosolic DNA sensing, type I interferon signaling, inflammatory pathways, and cGAS-STING-related responses.

#### Mouse infection experiments

Mouse infection experiments were performed to assess the contribution of ESAT-6 and CFP-10 to *in vivo* colonization, inflammation, and attenuation. C57BL/6 wild-type and *cGas^−/−^* mice were infected by intraperitoneal injection with 1 × 10^5^ CFU of the indicated *M. marinum* strains, with PBS as a control. At the indicated time point (120 h), the spleen, liver, and lung were collected for analysis of bacterial burden and inflammatory readouts. Tissue homogenates were plated to determine CFU per gram of tissue. For gene expression analysis, total RNA was extracted from tissues using the RNAprep Pure animal tissue total RNA kit (Tiangen, DP431, China), reverse-transcribed to cDNA, and analyzed by RT-qPCR. These experiments were used to compare wild-type, single-mutant, double-mutant, and complemented strains in different host genetic backgrounds.

#### Zebrafish infection experiments

Adult wild-type zebrafish (*Danio rerio*, AB strain) were maintained under standard conditions and infected by intraperitoneal injection with 1 × 10^5^ CFU of the indicated *M. marinum* strains; PBS/0.05% Tween-80 was used as a control. For survival analysis, fish were monitored daily for 24 days and analyzed by the Kaplan-Meier method. For bacterial burden analysis, fish were collected at 4 dpi, tissues were homogenized in PBS containing 0.05% Tween-80, serially diluted, and plated on 7H10 agar for CFU determination. For host response analysis, tissues were collected, snap-frozen, and processed for RT-qPCR to quantify gene expression.

#### Zebrafish immunization-challenge experiments

Zebrafish were subjected to a prime-boost-challenge regimen. Fish were immunized twice at a 7-day interval with PBS, wild-type *M. marinum*, or the Δ*esat-6*Δ*cfp-10* strain, followed by challenge with wild-type *M. marinum*. Survival was monitored daily after challenge and analyzed by the

Kaplan-Meier method. At 7 days post-challenge, fish were collected for RT-qPCR analysis of gene expression.

#### Statistics and reproducibility

All experiments were performed with at least three biological replicates and were repeated independently unless otherwise indicated. Data are presented as mean ± SD or mean ± SEM as specified in the figure legends. Statistical analyses were performed using GraphPad Prism 8.01. *P* values were calculated using two-tailed Student’s t-test for paired comparisons or one-way or two-way ANOVA for multiple comparisons. Error bars represent ± SEM. \**P* < 0.05; \*\**P* < 0.01; \*\*\**P* < 0.001; ns, not significant.

## Notes

### Competing Interest Statement

The authors have declared no competing interest.

## REFERENCES

1. Manzanillo, P.S., Shiloh, M.U., Portnoy, D.A., and Cox, J.S. (2012). *Mycobacterium tuberculosis* activates the DNA-dependent cytosolic surveillance pathway within macrophages. Cell Host Microbe 11, 469–480. 10.1016/j.chom.2012.03.007.

2. Houben, D., Demangel, C., van Ingen, J., Perez, J., Baldeon, L., Abdallah, A.M., Caleechurn, L., Bottai, D., van Zon, M., de Punder, K., et al. (2012). ESX-1-mediated translocation to the cytosol controls virulence of mycobacteria. Cell Microbiol 14, 1287–1298. 10.1111/j.1462-5822.2012.01799.x.

3. Simeone, R., Sayes, F., Song, O., Groschel, M.I., Brodin, P., Brosch, R., and Majlessi, L. (2015). Cytosolic access of *Mycobacterium tuberculosis*: critical impact of phagosomal acidification control and demonstration of occurrence *in vivo*. PLoS Pathog 11, e1004650. 10.1371/journal.ppat.1004650.

4. Conrad, W.H., Osman, M.M., Shanahan, J.K., Chu, F., Takaki, K.K., Cameron, J., Hopkinson-Woolley, D., Brosch, R., and Ramakrishnan, L. (2017). Mycobacterial ESX-1 secretion system mediates host cell lysis through bacterium contact-dependent gross membrane disruptions. Proc Natl Acad Sci U S A 114, 1371–1376. 10.1073/pnas.1620133114.

5. Watson, R.O., Bell, S.L., MacDuff, D.A., Kimmey, J.M., Diner, E.J., Olivas, J., Vance, R.E., Stallings, C.L., Virgin, H.W., and Cox, J.S. (2015). The Cytosolic Sensor cGAS Detects *Mycobacterium tuberculosis* DNA to Induce Type I Interferons and Activate Autophagy. Cell Host Microbe 17, 811–819. 10.1016/j.chom.2015.05.004.

6. Schorey, J.S., Vecchio, J., McManus, W.R., Ongalo, J., and Webber, K. (2024). Activation of host nucleic acid sensors by Mycobacterium: good for us or good for them? Crit Rev Microbiol 50, 224–240. 10.1080/1040841X.2023.2294904.

7. Chai, Q., Wang, L., Liu, C.H., and Ge, B. (2020). New insights into the evasion of host innate immunity by *Mycobacterium tuberculosis*. Cell Mol Immunol 17, 901–913. 10.1038/s41423-020-0502-z.

8. Hopfner, K.P., and Hornung, V. (2020). Molecular mechanisms and cellular functions of cGAS-STING signalling. Nat Rev Mol Cell Biol 21, 501–521. 10.1038/s41580-020-0244-x.

9. Collins, A.C., Cai, H., Li, T., Franco, L.H., Li, X.D., Nair, V.R., Scharn, C.R., Stamm, C.E., Levine, B., Chen, Z.J., and Shiloh, M.U. (2015). Cyclic GMP-AMP Synthase Is an Innate Immune DNA Sensor for *Mycobacterium tuberculosis*. Cell Host Microbe 17, 820–828. 10.1016/j.chom.2015.05.005.

10. Wassermann, R., Gulen, M.F., Sala, C., Perin, S.G., Lou, Y., Rybniker, J., Schmid-Burgk, J.L., Schmidt, T., Hornung, V., Cole, S.T., and Ablasser, A. (2015). *Mycobacterium tuberculosis* Differentially Activates cGAS- and Inflammasome-Dependent Intracellular Immune Responses through ESX-1. Cell Host Microbe 17, 799–810. 10.1016/j.chom.2015.05.003.

11. Donovan, M.L., Schultz, T.E., Duke, T.J., and Blumenthal, A. (2017). Type I Interferons in the Pathogenesis of Tuberculosis: Molecular Drivers and Immunological Consequences. Front Immunol 8, 1633. 10.3389/fimmu.2017.01633.

12. Moreira-Teixeira, L., Stimpson, P.J., Stavropoulos, E., Hadebe, S., Chakravarty, P., Ioannou, M., Aramburu, I.V., Herbert, E., Priestnall, S.L., Suarez-Bonnet, A., et al. (2020). Type I IFN exacerbates disease in tuberculosis-susceptible mice by inducing neutrophil-mediated lung inflammation and NETosis. Nat Commun 11, 5566. 10.1038/s41467-020-19412-6.

13. Zhang, L., Jiang, X., Pfau, D., Ling, Y., and Nathan, C.F. (2021). Type I interferon signaling mediates Mycobacterium tuberculosis-induced macrophage death. J Exp Med 218. 10.1084/jem.20200887.

14. Du, M., and Chen, Z.J. (2018). DNA-induced liquid phase condensation of cGAS activates innate immune signaling. Science 361, 704–709. 10.1126/science.aat1022.

15. Renshaw, P.S., Lightbody, K.L., Veverka, V., Muskett, F.W., Kelly, G., Frenkiel, T.A., Gordon, S.V., Hewinson, R.G., Burke, B., Norman, J., et al. (2005). Structure and function of the complex formed by the tuberculosis virulence factors CFP-10 and ESAT-6. EMBO J 24, 2491–2498. 10.1038/sj.emboj.7600732.

16. Aguilera, J., Karki, C.B., Li, L., Vazquez Reyes, S., Estevao, I., Grajeda, B.I., Zhang, Q., Arico, C.D., Ouellet, H., and Sun, J. (2020). N(alpha)-Acetylation of the virulence factor EsxA is required for mycobacterial cytosolic translocation and virulence. J Biol Chem 295, 5785–5794. 10.1074/jbc.RA119.012497.

17. Zhang, Q., Wang, D., Jiang, G., Liu, W., Deng, Q., Li, X., Qian, W., Ouellet, H., and Sun, J. (2016). EsxA membrane-permeabilizing activity plays a key role in mycobacterial cytosolic translocation and virulence: effects of single-residue mutations at glutamine 5. Sci Rep 6, 32618. 10.1038/srep32618.

18. Osman, M.M., Shanahan, J.K., Chu, F., Takaki, K.K., Pinckert, M.L., Pagan, A.J., Brosch, R., Conrad, W.H., and Ramakrishnan, L. (2022). The C terminus of the mycobacterium ESX-1 secretion system substrate ESAT-6 is required for phagosomal membrane damage and virulence. Proc Natl Acad Sci U S A 119, e2122161119. 10.1073/pnas.2122161119.

19. Champion, P.A. (2013). Disconnecting in vitro ESX-1 secretion from mycobacterial virulence. J Bacteriol 195, 5418–5420. 10.1128/JB.01145-13.

20. Abdallah, A.M., Weerdenburg, E.M., Guan, Q., Ummels, R., Borggreve, S., Adroub, S.A., Malas, T.B., Naeem, R., Zhang, H., Otto, T.D., et al. (2019). Integrated transcriptomic and proteomic analysis of pathogenic mycobacteria and their esx-1 mutants reveal secretion-dependent regulation of ESX-1 substrates and WhiB6 as a transcriptional regulator. PLoS One 14, e0211003. 10.1371/journal.pone.0211003.

21. Cronin, R.M., Ferrell, M.J., Cahir, C.W., Champion, M.M., and Champion, P.A. (2022). Proteo-genetic analysis reveals clear hierarchy of ESX-1 secretion in *Mycobacterium marinum*. Proc Natl Acad Sci U S A 119, e2123100119. 10.1073/pnas.2123100119.

22. Burts, M.L., Williams, W.A., DeBord, K., and Missiakas, D.M. (2005). EsxA and EsxB are secreted by an ESAT-6-like system that is required for the pathogenesis of Staphylococcus aureus infections. Proc Natl Acad Sci U S A 102, 1169–1174. 10.1073/pnas.0405620102.

23. Guo, Q., Bi, J., Li, M., Ge, W., Xu, Y., Fan, W., Wang, H., and Zhang, X. (2019). ESX Secretion-Associated Protein C From Mycobacterium tuberculosis Induces Macrophage Activation Through the Toll-Like Receptor-4/Mitogen-Activated Protein Kinase Signaling Pathway. Front Cell Infect Microbiol 9, 158. 10.3389/fcimb.2019.00158.

24. Guo, Y., Liu, K., Yang, X., Lv, Z., Zhao, K., Wang, X., Chu, Y., Li, J., and Huang, T. (2023). Multi-omics-based characterization of the influences of Mycobacterium tuberculosis virulence factors EsxB and PPE68 on host cells. Arch Microbiol 205, 230. 10.1007/s00203-023-03576-y.

25. Kennedy, G.M., Hooley, G.C., Champion, M.M., Mba Medie, F., and Champion, P.A. (2014). A novel ESX-1 locus reveals that surface-associated ESX-1 substrates mediate virulence in Mycobacterium marinum. J Bacteriol 196, 1877–1888. 10.1128/JB.01502-14.

26. Menon, A.R., Prest, R.J., Tobin, D.M., and Champion, P.A. (2025). *Mycobacterium marinum* as a model for understanding principles of mycobacterial pathogenesis. J Bacteriol 207, e0004725. 10.1128/jb.00047-25.

27. Chirakos, A.E., Balaram, A., Conrad, W., and Champion, P.A. (2020). Modeling Tubercular ESX-1 Secretion Using *Mycobacterium marinum*. Microbiol Mol Biol Rev 84, e00082–00019. 10.1128/MMBR.00082-19.

28. Groschel, M.I., Sayes, F., Simeone, R., Majlessi, L., and Brosch, R. (2016). ESX secretion systems: mycobacterial evolution to counter host immunity. Nat Rev Microbiol 14, 677–691. 10.1038/nrmicro.2016.131.

29. Chandra, P., Grigsby, S.J., and Philips, J.A. (2022). Immune evasion and provocation by Mycobacterium tuberculosis. Nat Rev Microbiol 20, 750–766. 10.1038/s41579-022-00763-4.

30. Shah, S., Bohsali, A., Ahlbrand, S.E., Srinivasan, L., Rathinam, V.A., Vogel, S.N., Fitzgerald, K.A., Sutterwala, F.S., and Briken, V. (2013). Cutting edge: *Mycobacterium tuberculosis* but not nonvirulent mycobacteria inhibits IFN-beta and AIM2 inflammasome-dependent IL-1beta production via its ESX-1 secretion system. J Immunol 191, 3514–3518. 10.4049/jimmunol.1301331.

31. Kranzusch, P.J., Lee, A.S., Berger, J.M., and Doudna, J.A. (2013). Structure of human cGAS reveals a conserved family of second-messenger enzymes in innate immunity. Cell Rep 3, 1362–1368. 10.1016/j.celrep.2013.05.008.

32. Zhou, W., Mohr, L., Maciejowski, J., and Kranzusch, P.J. (2021). cGAS phase separation inhibits TREX1-mediated DNA degradation and enhances cytosolic DNA sensing. Molecular Cell 81, 739–755.e737. 10.1016/j.molcel.2021.01.024.

33. Eaglesham, J.B., and Kranzusch, P.J. (2020). Conserved strategies for pathogen evasion of cGAS-STING immunity. Curr Opin Immunol 66, 27–34. 10.1016/j.coi.2020.04.002.

34. Dey, R.J., Dey, B., Zheng, Y., Cheung, L.S., Zhou, J., Sayre, D., Kumar, P., Guo, H., Lamichhane, G., Sintim, H.O., and Bishai, W.R. (2017). Inhibition of innate immune cytosolic surveillance by an *M. tuberculosis* phosphodiesterase. Nat Chem Biol 13, 210–217. 10.1038/nchembio.2254.

35. Dey, B., Dey, R.J., Cheung, L.S., Pokkali, S., Guo, H., Lee, J.H., and Bishai, W.R. (2015). A bacterial cyclic dinucleotide activates the cytosolic surveillance pathway and mediates innate resistance to tuberculosis. Nat Med 21, 401–406. 10.1038/nm.3813.

36. Murdoch, C.C., and Skaar, E.P. (2022). Nutritional immunity: the battle for nutrient metals at the host-pathogen interface. Nat Rev Microbiol 20, 657–670. 10.1038/s41579-022-00745-6.

37. Zhou, W., Mohr, L., Maciejowski, J., and Kranzusch, P.J. (2021). cGAS phase separation inhibits TREX1-mediated DNA degradation and enhances cytosolic DNA sensing. Mol Cell 81, 739–755 e737. 10.1016/j.molcel.2021.01.024.

38. Sun, Y., Zhang, W., Dong, C., and Xiong, S. (2020). *Mycobacterium tuberculosis* MmsA (Rv0753c) Interacts with STING and Blunts the Type I Interferon Response. mBio 11, e03254–03219. 10.1128/mBio.03254-19.

39. Nieuwenhuizen, N.E., and Kaufmann, S.H.E. (2018). Next-Generation Vaccines Based on Bacille Calmette-Guérin. Front Immunol 9, 121. 10.3389/fimmu.2018.00121.

40. Wang, C., Guan, Y., Lv, M., Zhang, R., Guo, Z., Wei, X., Du, X., Yang, J., Li, T., Wan, Y., et al. (2018). Manganese Increases the Sensitivity of the cGAS-STING Pathway for Double-Stranded DNA and Is Required for the Host Defense against DNA Viruses. Immunity 48, 675–687.e677. 10.1016/j.immuni.2018.03.017.

41. Song, L., Xu, L., Zhang, P., Li, S., Qu, Y., Zhao, Y., Shi, Z., Ma, R., Li, Y., Chen, Y., et al. (2025). A Dual-Targeting T6SS DNase Drives Bacterial Antagonism and Eukaryotic Apoptosis via the cGAS-STING-TNF Axis. Adv Sci (Weinh) 12, e2504086. 10.1002/advs.202504086.

42. Zhu, L., Zuo, Y., Cui, R., Fu, P., Liu, Y., Wang, Z., He, X., Yu, D., Wei, Z., Li, S., et al. (2026). Interkingdom sensing of fungal tyrosol promotes bacterial antifungal T6SS activity in the murine gut. Nat Microbiol 11, 240–255. 10.1038/s41564-025-02208-z.

43. Wang, X., Li, Y., Wang, X., Zhang, W., Liu, M., Hao, X., Chen, S., Chang, T., Wu, C., Hao, C., et al. (2026). Manganese activates the CBASS immunity to protect bacteria from phage infection. mBio 17, e0275825. 10.1128/mbio.02758-25.

44. Wang, T., Si, M., Song, Y., Zhu, W., Gao, F., Wang, Y., Zhang, L., Zhang, W., Wei, G., Luo, Z.Q., and Shen, X. (2015). Type VI Secretion System Transports Zn2+ to Combat Multiple Stresses and Host Immunity. PLoS Pathog 11, e1005020. 10.1371/journal.ppat.1005020.

45. Zhu, L., Xu, L., Wang, C., Li, C., Li, M., Liu, Q., Wang, X., Yang, W., Pan, D., Hu, L., et al. (2021). T6SS translocates a micropeptide to suppress STING-mediated innate immunity by sequestering manganese. Proc Natl Acad Sci U S A 118, e2103526118. 10.1073/pnas.2103526118.

