## Supplemental Figures for "ESX-1 substrates ESAT-6 and CFP-10 limit cGAS-STING activation via DNA and Zn^2+^ restriction"

**SUPPLEMENTAL FIGURE LEGENDS**

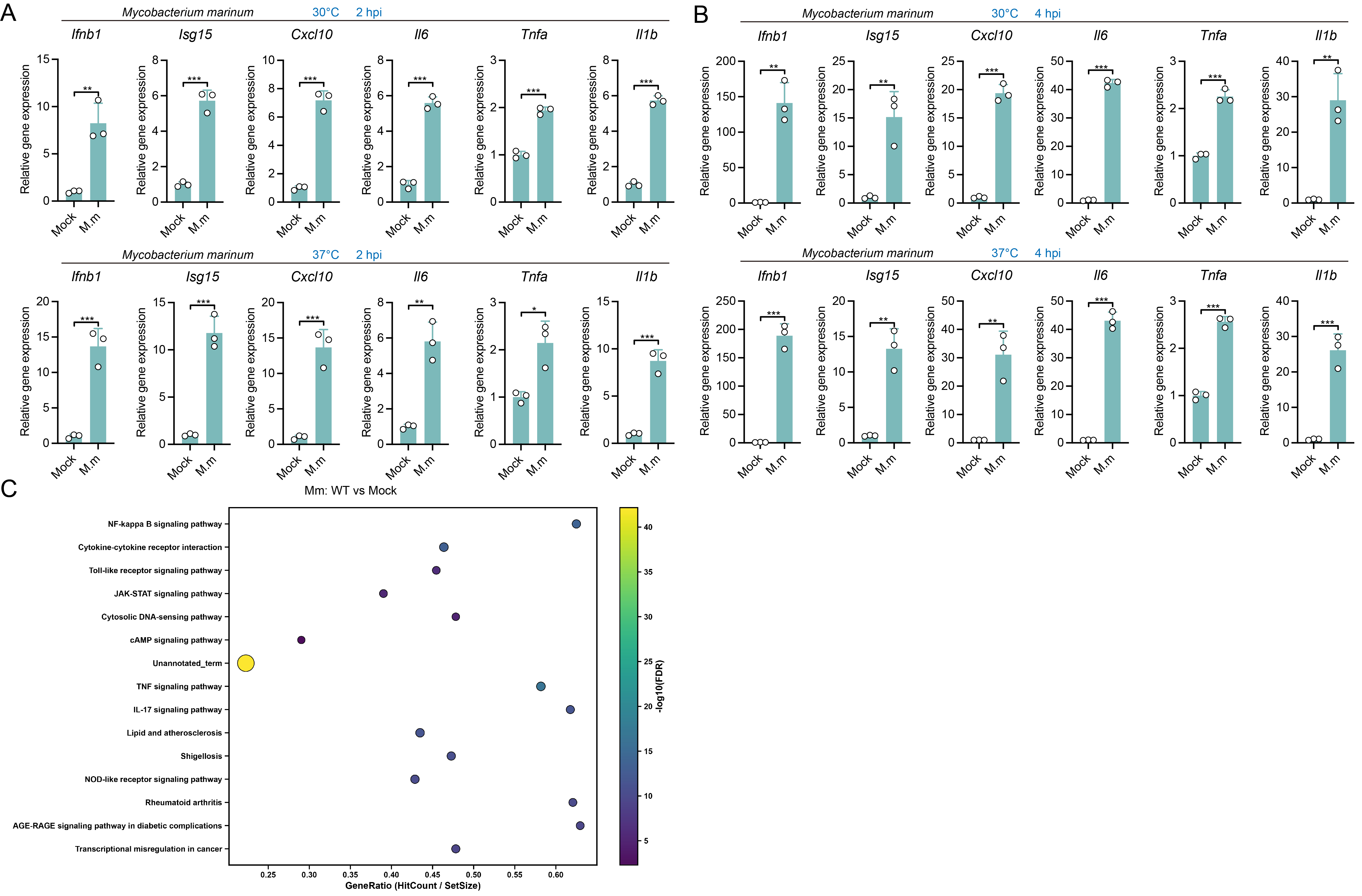

**Figure S1. *Mycobacterium marinum* infection induces innate immune gene expression in macrophages.**

(A) RT-qPCR analysis of gene expression in primary macrophages infected with *M. marinum* (MOI = 50) or mock-treated controls at 30°C or 37°C for 2 h. Gene expression was normalized to the indicated internal control and presented as relative expression values. n = 3.

(B) RT-qPCR analysis of gene expression in primary macrophages infected with *M. marinum* (MOI = 50) or mock-treated controls at 30°C or 37°C for 4 h. Gene expression was normalized to the indicated internal control and presented as relative expression values. n = 3.

(C) Pathway enrichment analysis of differentially expressed genes identified by RNA-seq in primary macrophages infected with WT *M. marinum* (MOI = 20, 4 h) relative to mock-treated controls. n = 3.

*P* values were calculated using two-tailed Student’s t-test for paired comparisons or one-way or two-way ANOVA for multiple comparisons.

Error bars represent ± SEM. **P* < 0.05; ***P* < 0.01; ****P* < 0.001; ns, not significant.

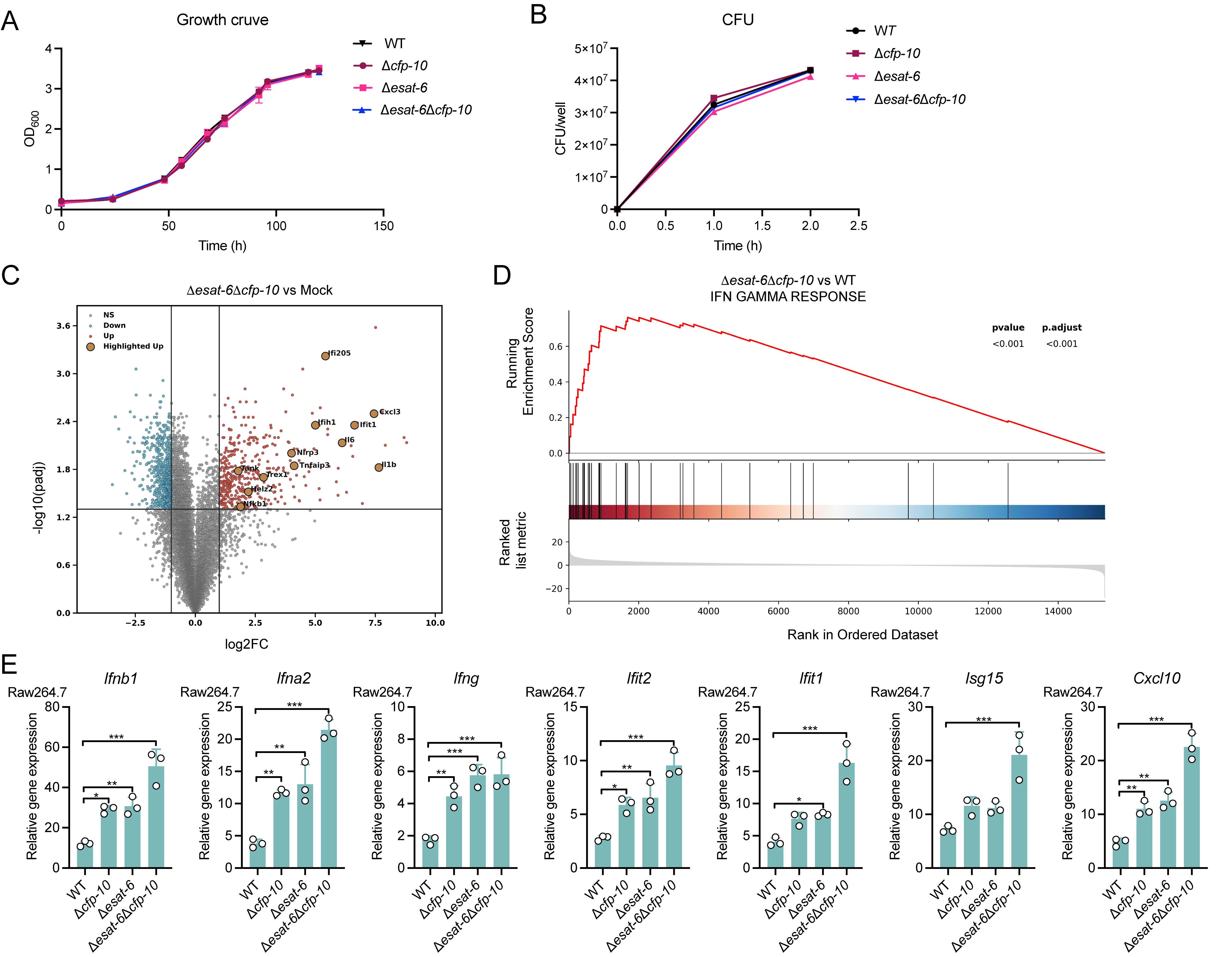

**Figure S2. Supporting analyses for ESAT-6/CFP-10-mediated restraint of interferon-associated responses during mycobacterial infection.**

(A) In vitro growth curves of WT, Δ*cfp-10*, Δ*esat-6*, and Δ*esat-6*Δ*cfp-10* *M. marinum* strains measured by OD_600_ over the indicated time course. n = 3.

(B) Quantification of intracellular bacterial burdens in macrophages infected with WT, Δ*cfp-10*, Δ*esat-6*, and Δ*esat-6*Δ*esx* strains (MOI =10) at the indicated early infection time points, expressed as CFU per well. n = 3.

(C) Volcano plot showing differentially expressed genes identified by RNA-seq in primary macrophages infected with the Δ*esat-6*Δ*cfp-10* strain (MOI = 20, 4 h) relative to mock-treated controls. Highlighted genes are indicated. n = 3.

(D) Gene set enrichment analysis (GSEA) performed on ranked transcriptomic data comparing Δ*esat-6*Δ*cfp-10*-infected macrophages with WT-infected macrophages (MOI = 20, 4 h) using the IFN GAMMA RESPONSE gene set. n = 3.

(E) RT-qPCR analysis of gene expression in Raw264.7 macrophages infected with WT, Δ*cfp-10*, Δ*esat-6*, or Δ*esat-6*Δ*cfp-10* strains (MOI =20, 4 h). Gene expression was normalized to the indicated internal control and presented as relative expression values. n = 3.

*P*  values calculated using two-tailed Student’s t-test for paired comparisons or one-way or two way ANOVA for multiple comparisons.

Error bars represent ± SEM. **P* < 0.05; ***P* < 0.01; ****P* < 0.001; ns, not significant.

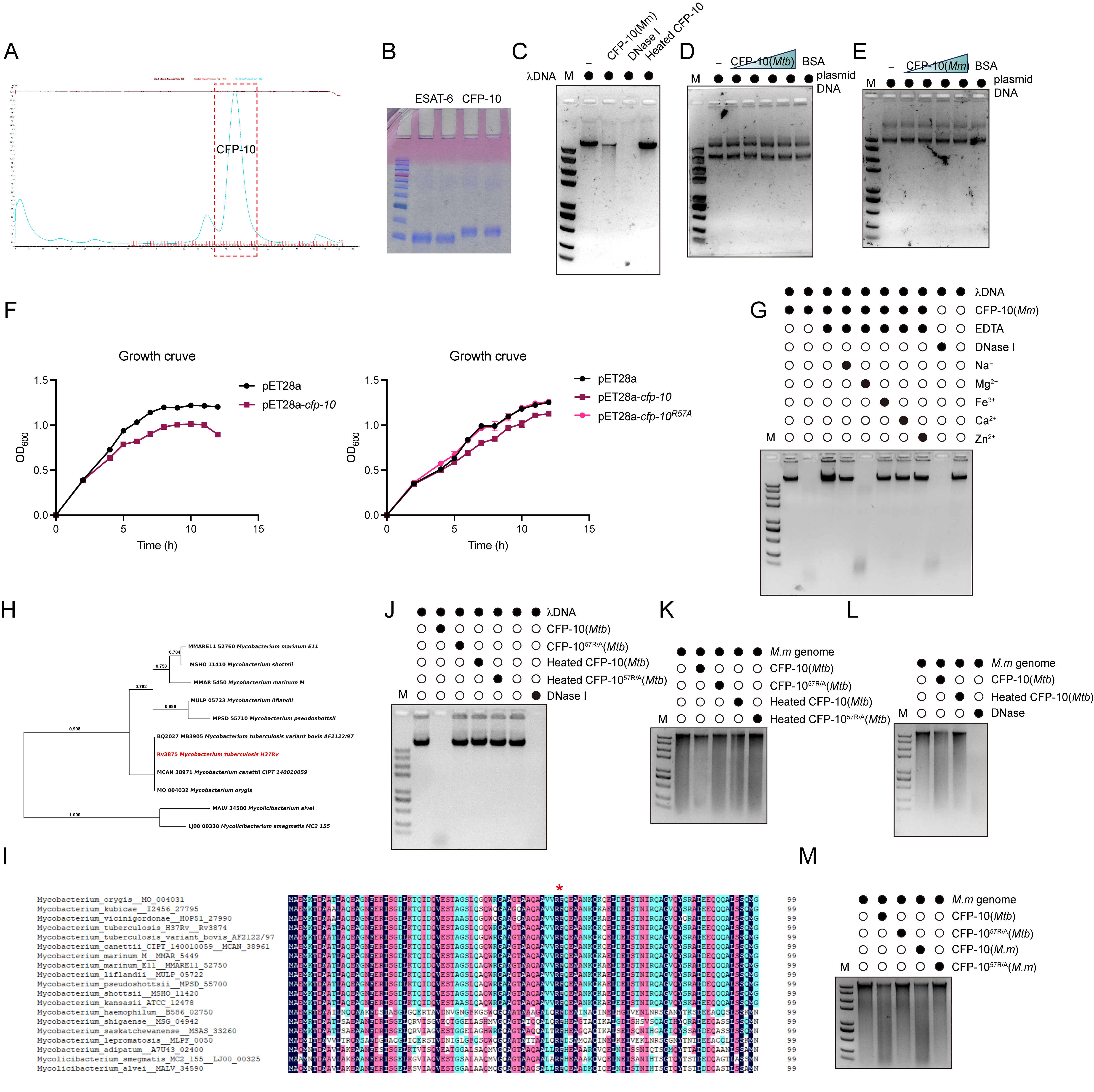

**Figure S3. Biochemical analyses of CFP-10 as a metal-dependent DNase.**

(A) Size-exclusion chromatography profile of recombinant CFP-10, with the indicated elution peak highlighted.

(B) Coomassie-stained SDS-PAGE analysis of purified recombinant Rv3875 and Rv3874 proteins.

(C) *In vitro* DNA degradation assay in which linear λDNA (20 ng) was incubated with recombinant *M. marinum* CFP-10 (10 μM), DNase I (1 U/μL), or heat-inactivated CFP-10 (10 μM) and analyzed by agarose gel electrophoresis.

(D) Plasmid DNA processing assay using increasing amounts of recombinant *M. tuberculosis* CFP-10 (5, 9, 11, 14 μM) or BSA (8 μM), followed by agarose gel electrophoresis.

(E) Plasmid DNA processing assay using increasing amounts of recombinant *M. marinum* CFP-10 (5, 9, 11, 14 μM) or BSA (8 μM), followed by agarose gel electrophoresis.

(F) Growth curves of *E. coli* expressing empty vector, *cfp-10*, or *cfp-10*^R57A^, measured by OD_600_ over the indicated time course. Cultures were induced with 0.5 mM IPTG when the OD_600_ reached approximately 0.3. n = 3.

(G) Metal-dependence assay in which λDNA (20 ng) was incubated with recombinant *M. marinum* CFP-10 (10 μM) in the presence of EDTA (25 mM) or the indicated metal ions (4 mM Na⁺, 4 mM Mg^2+^, 1 mM Fe^3+^, 4 mM Ca^2+^, and 1 mM Zn^2+^), followed by agarose gel analysis.

(H) Phylogenetic analysis of CFP-10 homologs from the indicated mycobacterial species.

(I) Multiple sequence alignment of CFP-10 homologs from the indicated mycobacterial species, with the conserved R57-containing region marked.

(J) DNA degradation assay comparing wild-type *M. tuberculosis* CFP-10 (10 μM), CFP-10^R57A^ (10 μM), heat-inactivated CFP-10 (10 μM), and heat-inactivated CFP-10^R57A^ (10 μM), with DNase I (1 U/μL) included as a control.

(K) Genomic DNA degradation assay in which *M. marinum* genomic DNA (100 ng) was incubated with recombinant M. tuberculosis CFP-10 (10 μM), CFP-10^R57A^ (10 μM), or heat-inactivated CFP-10 (10 μM) and analyzed by agarose gel electrophoresis.

(L) Genomic DNA (100 ng) degradation assay in which *M. marinum* genomic DNA was incubated with recombinant M. tuberculosis CFP-10 (10 μM), heat-inactivated CFP-10 (10 μM), or DNase I (1 U/μL) and analyzed by agarose gel electrophoresis.

(M) Genomic DNA degradation assay comparing *M. tuberculosis* CFP-10 (10 μM), CFP-10^R57A^ (10 μM), *M. marinum* CFP-10 (10 μM), and *M. marinum* CFP-10^R57A^ (10 μM), followed by agarose gel electrophoresis. Representative of 3 independent experiments.

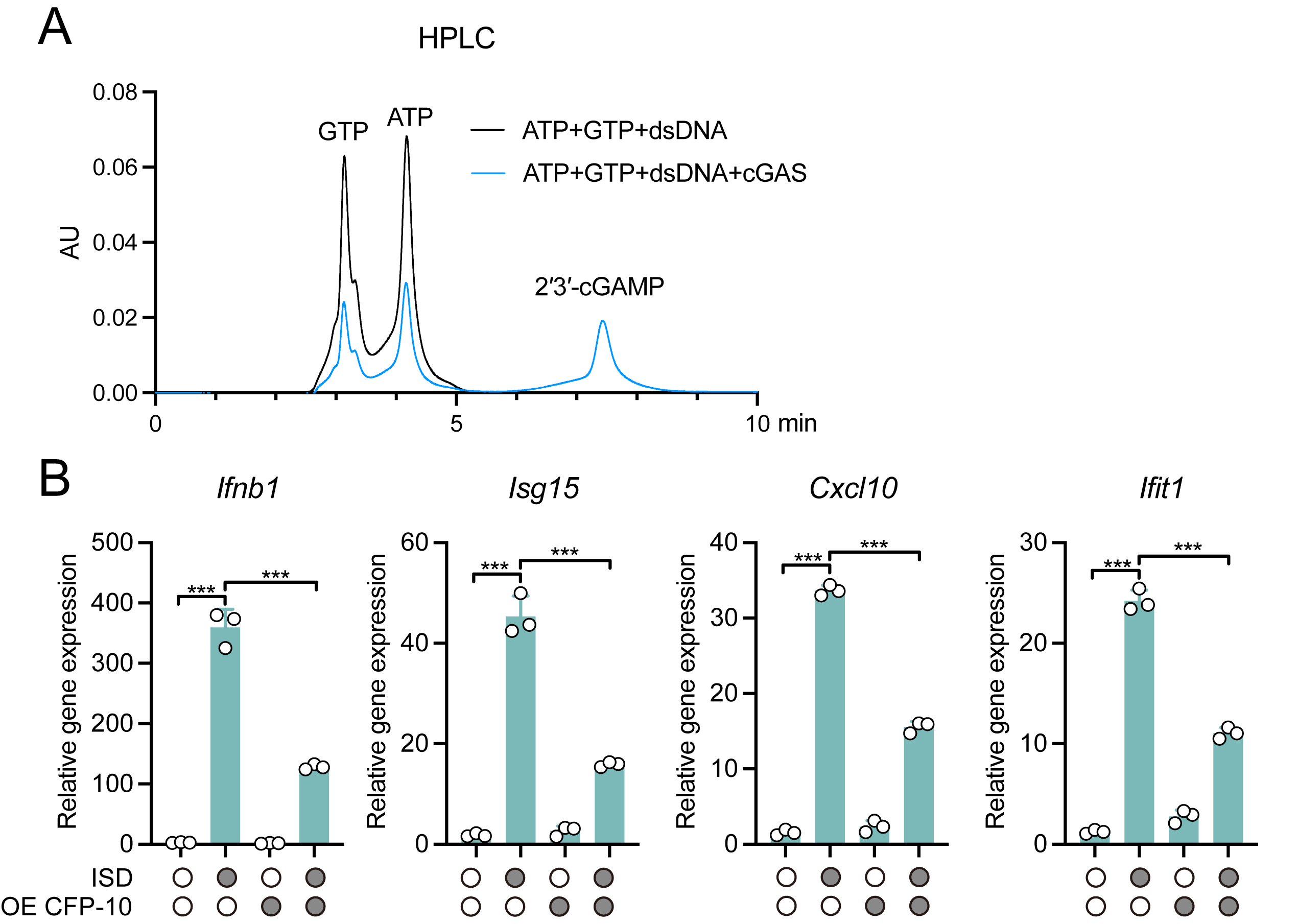

**Figure S4. CFP-10-mediated suppression of DNA-triggered cGAS activation.**

(A) HPLC analysis of nucleotide products from an in vitro reaction containing ATP (0.2 mM), GTP (0.2 mM), and dsDNA (10^−2^ mg/mL) in the presence or absence of recombinant cGAS (10 μM). Peaks corresponding to GTP, ATP, and 2′3′-cGAMP are indicated. Representative of 3 independent experiments.

(B) RT-qPCR analysis of gene expression in PMs transfected with ISD for 4 h and expressing CFP-10, as indicated. n = 3.

*P* values were calculated using two-tailed Student’s t-test for paired comparisons or one-way or two-way ANOVA for multiple comparisons.

Error bars represent ± SEM. ****P* < 0.001.

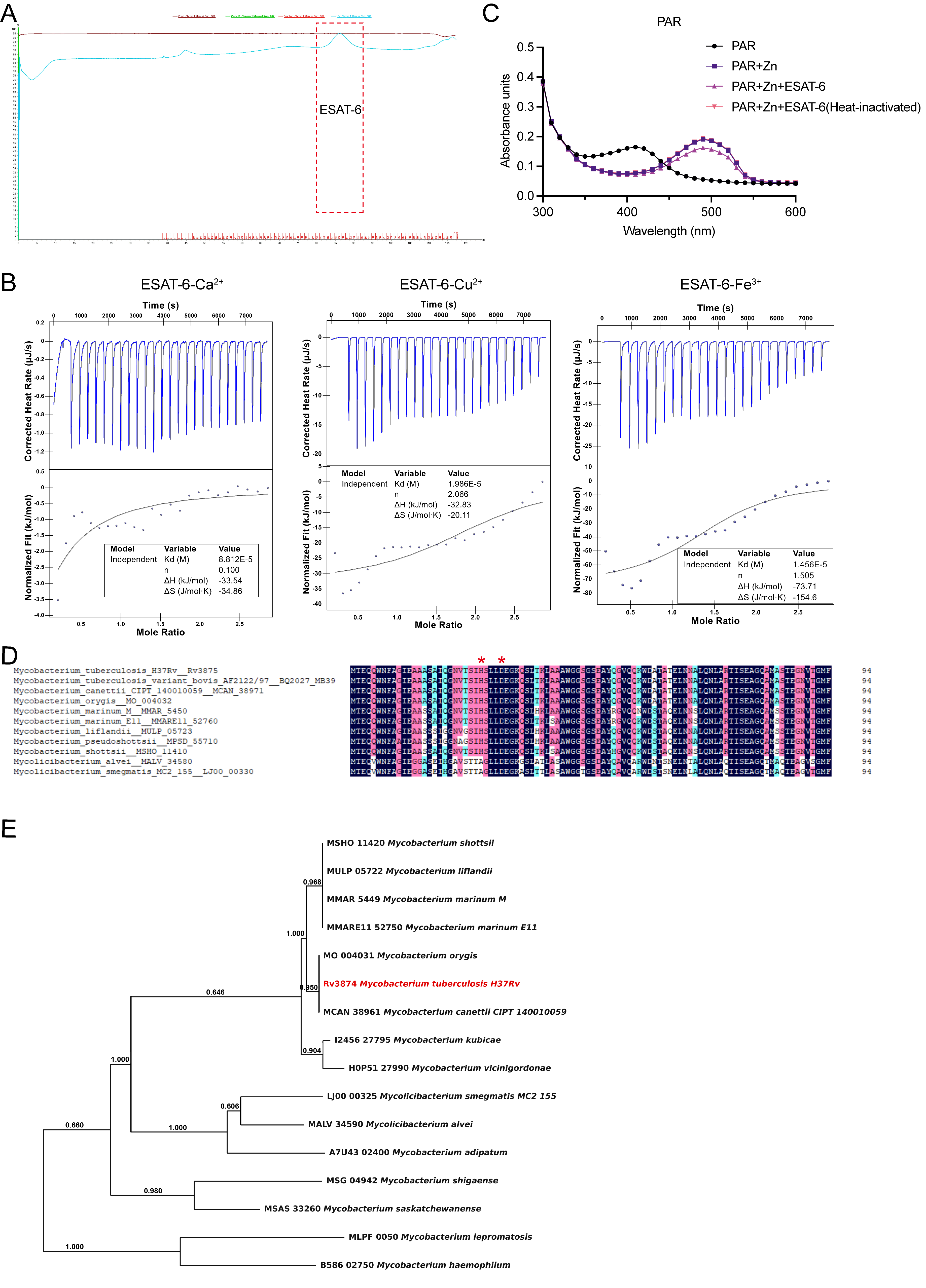

**Figure S5. Biochemical characterization of ESAT-6 Zn^2+^-binding ability.**

(A) Size-exclusion chromatography profile of recombinant ESAT-6, with the indicated elution peak highlighted.

(B) Isothermal titration calorimetry (ITC) analysis of recombinant ESAT-6 (25 μM) with Ca^2+^ (1 mM), Cu^2+^ (1 mM), or Fe^3+^ (1 mM). Upper panels show raw heat change upon sequential injections; lower panels show fitted binding isotherms.

(C) PAR competition assay performed with PAR, Zn^2+^, wild-type ESAT-6, or heat-inactivated ESAT-6, with absorbance spectra recorded across the indicated wavelengths.

(D) Multiple sequence alignment of ESAT-6 homologs from the indicated mycobacterial species, with the conserved H26/D30-containing region marked.

(E) Phylogenetic analysis of ESAT-6 homologs from the indicated mycobacterial species.

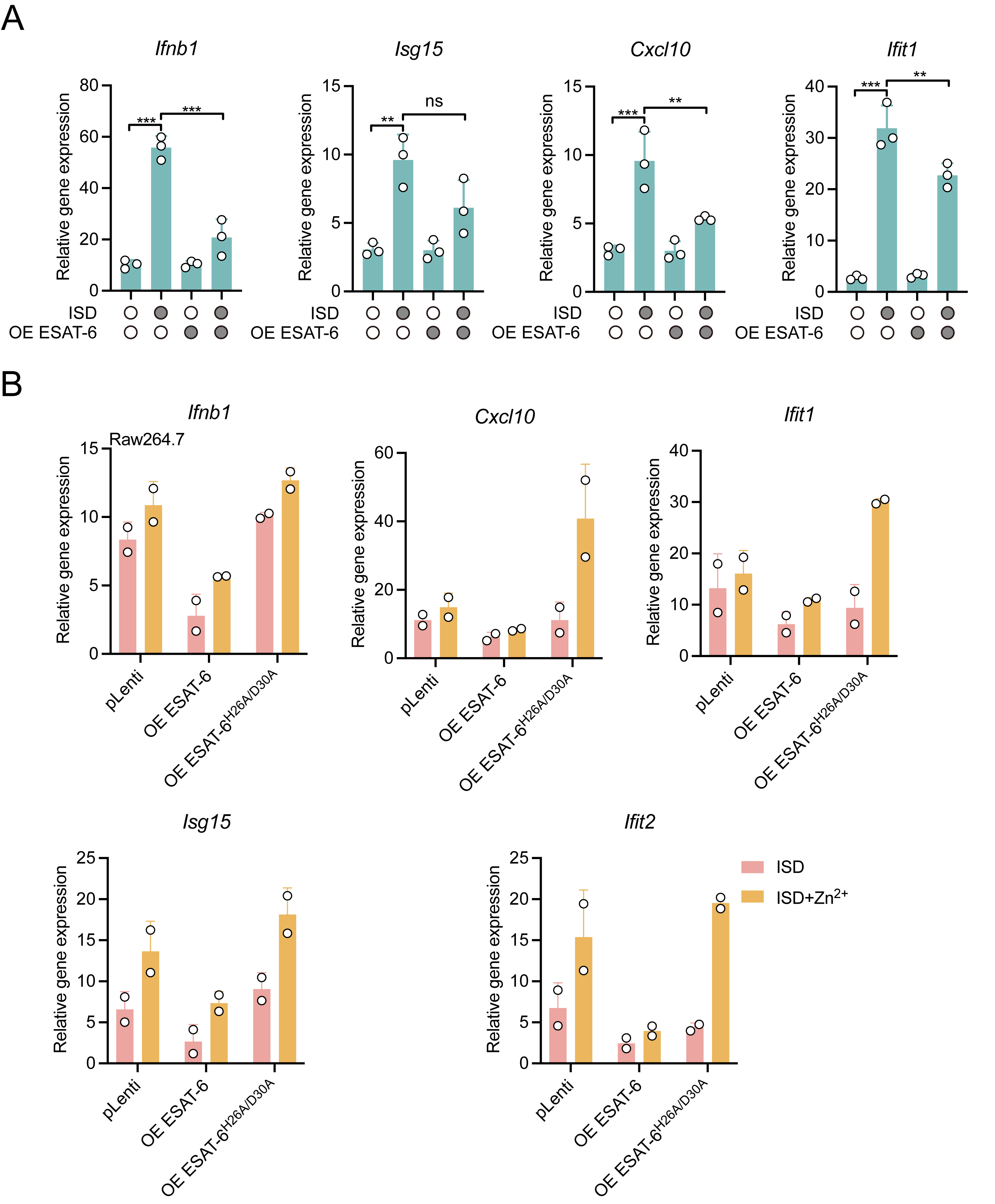

**Figure S6. ESAT-6 suppresses ISD-induced innate immune gene expression in a Zn^2+^-dependent manner.**

(A) RT-qPCR analysis of gene expression in cells transfected with ISD for 4 h and expressing ESAT-6, as indicated. n = 3.

(B) RT-qPCR analysis of gene expression in RAW 264.7 cells transfected with ISD for 4 h with or without Zn supplementation and expressing vector control, wild-type ESAT-6, or ESAT-6^H26A/D30A^. n = 2.

*P* values were calculated using two-tailed Student’s t-test for paired comparisons or one-way or two-way ANOVA for multiple comparisons.

Error bars represent ± SEM. **P* < 0.05; ***P* < 0.01; ****P* < 0.001; ns, not significant.

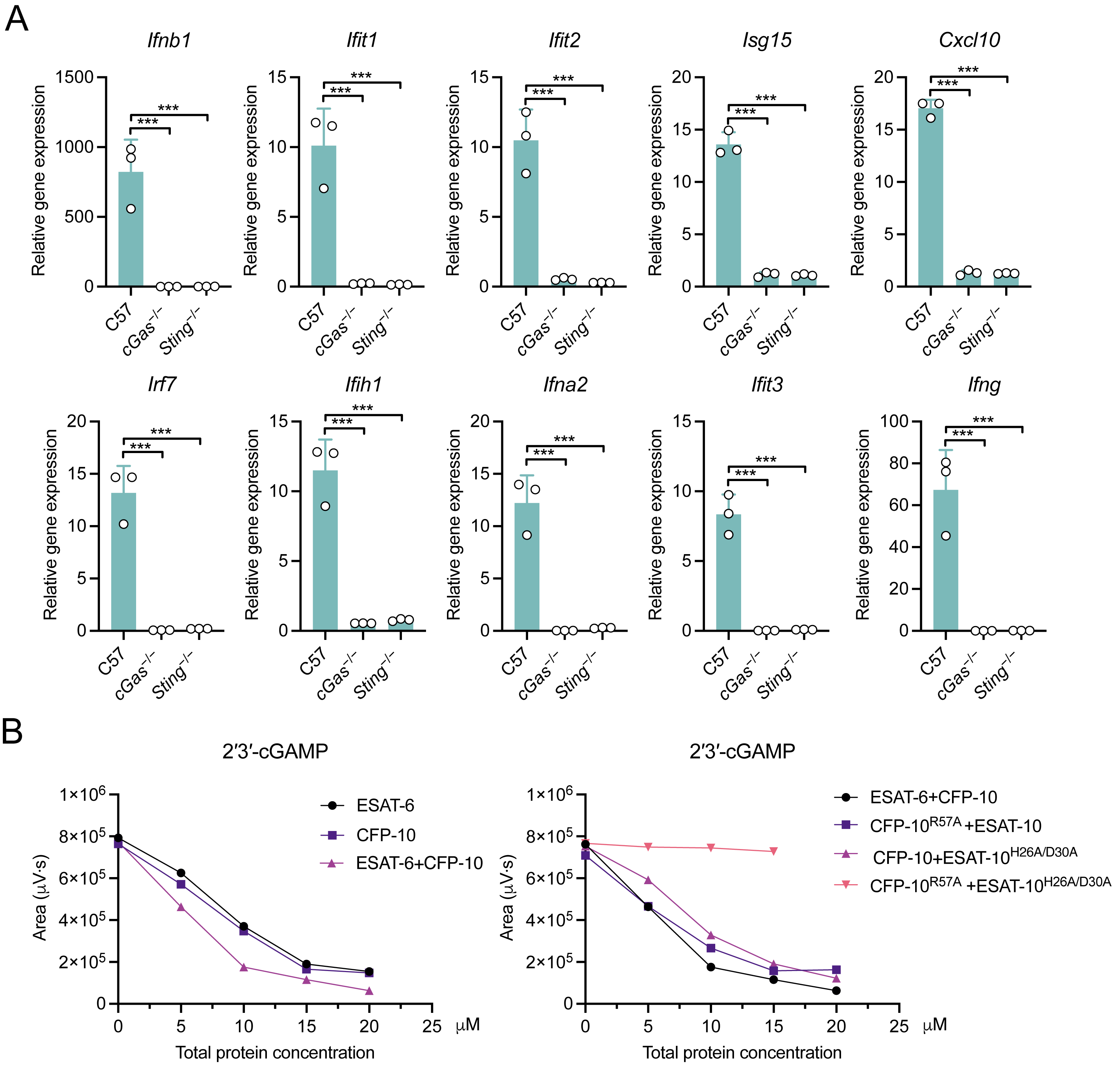

**Figure S7. cGAS-STING dependence of WT mycobacterial responses and biochemical cooperation between ESAT-6 and CFP-10.**

(A) RT-qPCR analysis of gene expression in C57, *cGas^−/−^*, and *Sting^−/−^* macrophages infected with WT *M. marinum* (MOI = 20, 4 h). n = 3.

(B) Quantification of 2′3′-cGAMP production by HPLC in reconstituted in vitro cGAS assays containing ESAT-6, CFP-10, ESAT-6 plus CFP-10, or combinations containing CFP-10^R57A^ and/or ESAT-6^H26A/D30A^ as indicated.

*P* values were calculated using two-tailed Student’s t-test for paired comparisons or one-way or two-way ANOVA for multiple comparisons.

Error bars represent ± SEM. **P* < 0.05; ***P* < 0.01; ****P* < 0.001; ns, not significant.

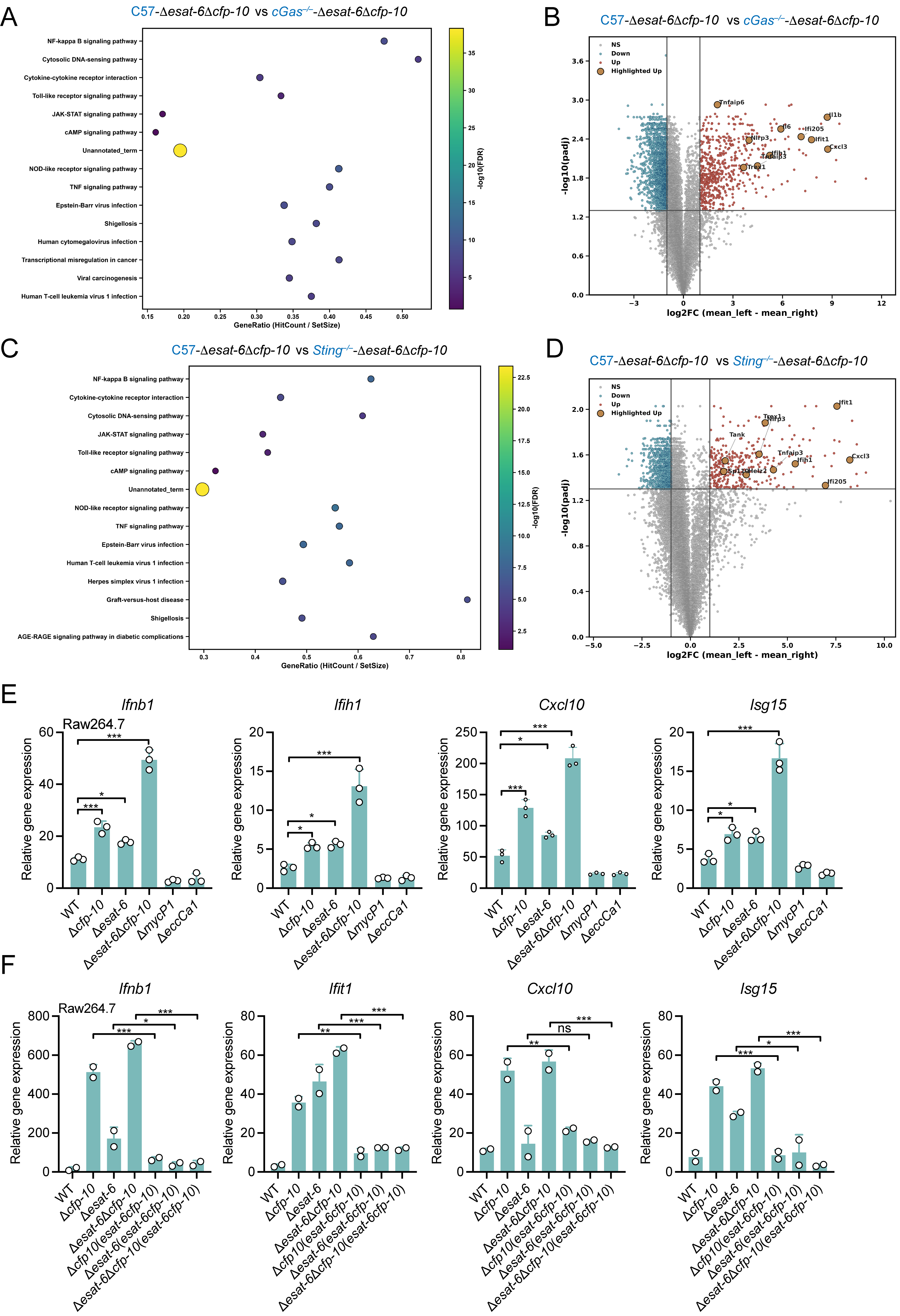

**Figure S8. Transcriptomic and Raw264.7 analyses of ESAT-6/CFP-10-dependent immune regulation.**

(A) Pathway enrichment analysis of genes differentially expressed between C57 and *cGas^−/−^* macrophages after Δ*esat-6*Δ*cfp-10* infection (MOI = 20, 4 h). n = 3 independent biological replicates per group.

(B) Volcano plot showing differentially expressed genes between C57 and *cGas^−/−^* macrophages after Δ*esat-6*Δ*cfp-10* infection (MOI = 20, 4 h). Highlighted genes are indicated. n = 3 independent biological replicates per group.

(C) Pathway enrichment analysis of genes differentially expressed between C57 and *Sting^−/−^* macrophages afterΔ*esat-6*Δ*cfp-10* infection (MOI = 20, 4 h). n = 3 independent biological replicates per group.

(D) Volcano plot showing differentially expressed genes between C57 and *Sting^−/−^* macrophages after Δ*esat-6*Δ*cfp-10* infection. Highlighted genes are indicated (MOI = 20, 4 h). n = 3 independent biological replicates per group.

(E) RT-qPCR analysis of gene expression in Raw264.7 macrophages infected with WT, Δ*cfp-10*, Δ*esat-6*, Δ*esat-6*Δ*cfp-10*, Δ*mycP1*, or Δ*eccCa1* strains (MOI = 20, 4 h). Gene expression was normalized to the indicated internal control and presented as relative expression values. n = 3.

(F) RT-qPCR analysis of gene expression in Raw264.7 macrophages infected with WT, deletion, and complementation strains carrying wild-type or mutant esat-6/cfp-10 alleles (MOI = 20, 4 h). Gene expression was normalized to the indicated internal control and presented as relative expression values. n = 2 independent biological replicates per group.

*P*  values calculated using two-tailed Student’s t-test for paired comparisons or one-way or two way ANOVA for multiple comparisons.

Error bars represent ± SEM. **P* < 0.05; ***P* < 0.01; ****P* < 0.001; ns, not significant.

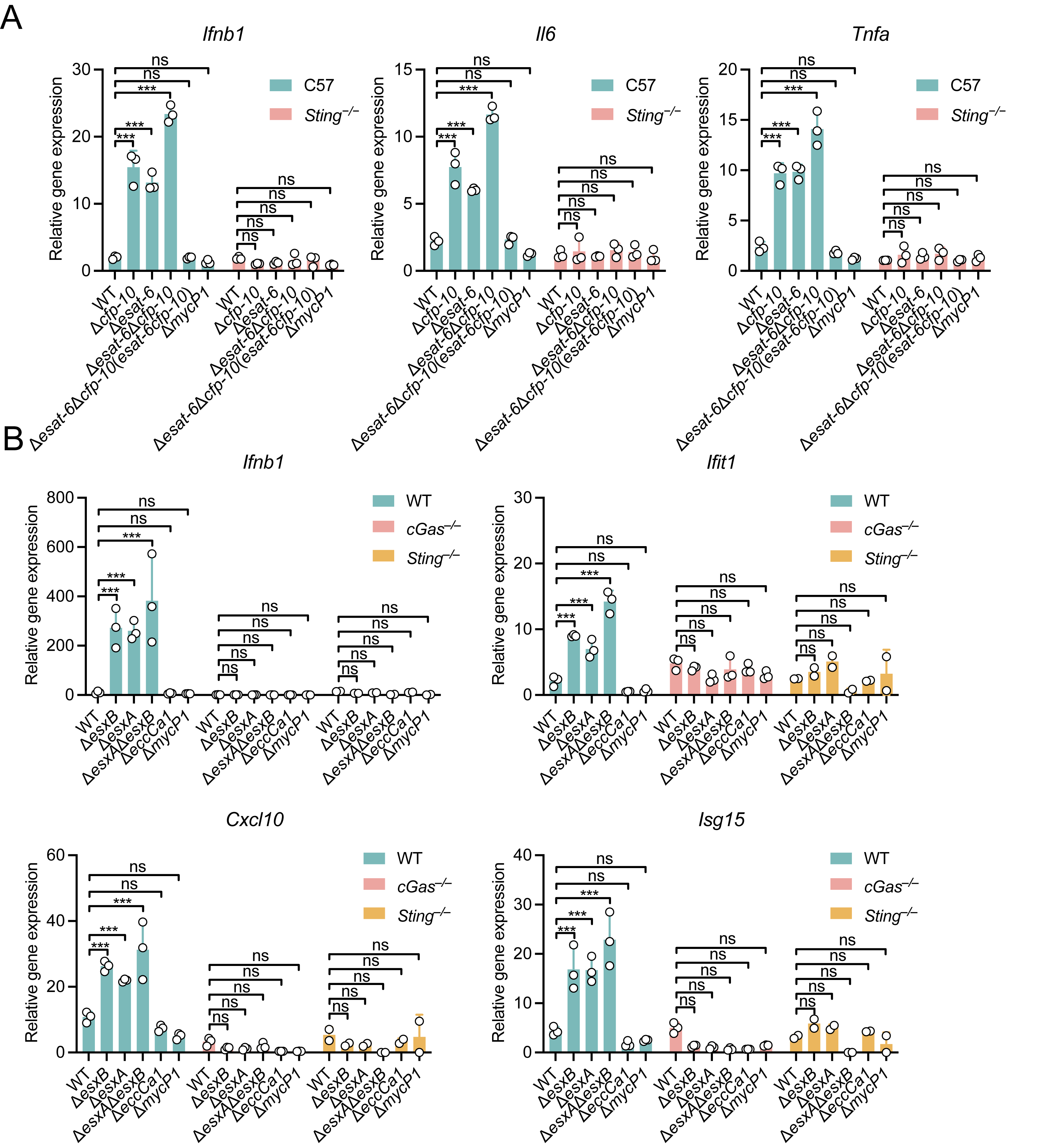

**Figure S9. Figure S9. Loss of cGAS or STING abolishes enhanced innate immune activation induced by esat-6/cfp-10 mutant strains.**

(A) RT-qPCR analysis of gene expression in C57 and *Sting^−/−^* macrophages infected with the indicated WT, deletion, complementation, or Δ*mycP1* strains (MOI = 20, 4 h). Gene expression was normalized to the indicated internal control and presented as relative expression values. n = 3.

(B) RT-qPCR analysis of gene expression in WT, *cGas^−/−^*, and *Sting^−/−^* macrophages infected with the indicated WT, deletion, complementation, or Δ*mycP1* strains (MOI = 20, 4 h). Gene expression was normalized to the indicated internal control and presented as relative expression values. n = 3.

*P*  values calculated using two-tailed Student’s t-test for paired comparisons or one-way or two way ANOVA for multiple comparisons.

Error bars represent ± SEM. ****P* < 0.001; ns, not significant.

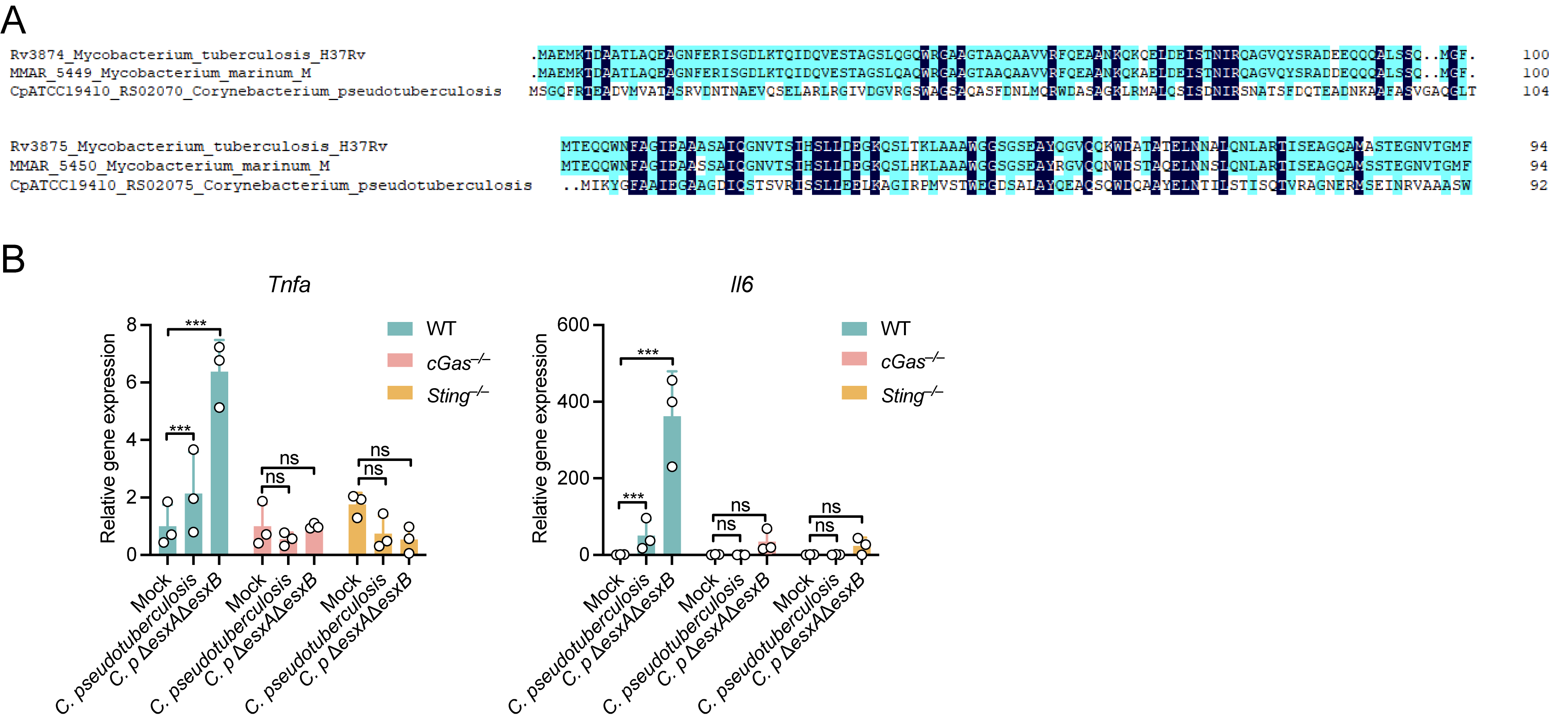

**Figure S10. cGAS-STING-dependent responses to *Corynebacterium pseudotuberculosis*.**

(A) Multiple sequence alignment of CFP-10-like and ESAT-6-like proteins from *M. tuberculosis* H37Rv, *M. marinum,* and *Corynebacterium pseudotuberculosis*.

(B) RT-qPCR analysis of gene expression in WT, *cGas^−/−^*, and *Sting^−/−^* macrophages infected with C. pseudotuberculosis or the corresponding Δ*esat-6*Δ*cfp-10* strain (MOI = 10, 4 h). Gene expression was normalized to the indicated internal control and presented as relative expression values. n = 3–4.

*P*  values calculated using two-tailed Student’s t-test for paired comparisons or one-way or two way ANOVA for multiple comparisons.

Error bars represent ± SEM. ****P* < 0.001; ns, not significant.
