## Supplemental Tables for "ESX-1 substrates ESAT-6 and CFP-10 limit cGAS-STING activation via DNA and Zn^2+^ restriction"

**Table S1. Bacterial strains used in this study.**

| **Strain** | **Relevant characteristics** | **Reference** |
| --- | --- | --- |
| ***E. coli*** |  |  |
| DH5α | FΦ80Δ*lacZ*ΔM15/Δ(*lacZYA-argF*)*U169recA1 endA1 hsdR17* | Novagen |
| BL21 (DE3) | Host for expression vector pGEX6P-1 | Novagen |
| ***M. marinum*** |  |  |
| WT | Wild-type does not carry antibiotic resistance | This study |
| Δ*cfp-10* (Mm) | *cfp-10* (Mm) gene deleted in *M. marinum* | This study |
| Δ*cfp-10* (Mm) (pMV261-*cfp-10* (Mm)) | Δ*cfp-10* (Mm) containing pMV261-*cfp-10* (Mm), Km^r^ | This study |
| Δ*cfp-10* (Mm) (pMV261-*cfp-10*^R57A^ (Mm)) | Δ*cfp-10* (Mm) containing pMV261-*cfp-10*^R57A^ (Mm), Km^r^ | This study |
| Δ*cfp-10* (Mm) (pMV261-*cfp-10* (Mtb)) | Δ*cfp-10* (Mm) containing pMV261-*cfp-10* (Mtb), Km^r^ | This study |
| Δ*cfp-10* (Mm) (pMV261-*cfp-10*^R57A^ (Mtb)) | Δ*cfp-10* (Mm) containing pMV261-*cfp-10*^R57A^ (Mtb), Km^r^ | This study |
| Δ*esat-6* (Mm) | *esat-6* (Mm) gene deleted in *M. marinum* | This study |
| Δ*esat-6* (Mm) (pMV261-*esat-6* (Mm)) | Δ*esat-6* (Mm) containing pMV261-*esat-6* (Mm), Km^r^ | This study |
| Δ*esat-6* (Mm) (pMV261-*esat-6*^H26A/D30A^ (Mm)) | Δ*esat-6* (Mm) containing pMV261-*esat-6*^H26A/D30A^ (Mm), Km^r^ | This study |
| Δ*esat-6* (Mm) (pMV261-*esat-6* (Mtb)) | Δ*esat-6* (Mm) containing pMV261-*esat-6* (Mtb), Km^r^ | This study |
| Δ*esat-6* (Mm) (pMV261-*esat-6*^H26A/D30A^ (Mtb)) | Δ*esat-6* (Mm) containing pMV261-*esat-6*^H26A/D30A^ (Mtb), Km^r^ | This study |
| Δ*esat-6*Δ*cfp-10* (Mm) | *esat-6* (Mm) and *cfp-10* (Mm) genes deleted in *M. marinum* | This study |
| Δ*esat-6*Δ*cfp-10* (Mm) (pMV261-Δ*esat-6*Δ*cfp-10* (Mm)) | Δ*esat-6*Δ*cfp-10* (Mm) containing pMV261-Δ*esat-6*Δ*cfp-10* (Mm), Km^r^ | This study |
| Δ*esat-6*Δ*cfp-10* (Mm) (pMV261-*cfp-10-esat-6* (Mtb)) | Δ*esat-6*Δ*cfp-10* (Mm) containing pMV261-*cfp-10-esat-6* (Mtb), Km^r^ | This study |
| Δ*mycP1* | *mycP1* gene deleted in *M. marinum* | This study |
| Δ*eccCA1* | *eccCA1* gene deleted in *M. marinum* | This study |

*Km, kanamycin; the working concentrations were 25 μg/mL.

**Table S2. Plasmids used in this study.**

| **Plasmid** | **Relevant characteristics** | **Reference** |
| --- | --- | --- |
| pMV261 | *E. coli*–*Mycobacterium* shuttle expression vector, pAL5000 replicon, Km^r^ | (Seniya *et al.*, 2020) |
| pMV261-*cfp-10* (Mm) | *cfp-10* (Mm) under the control of the promoter in plasmid pMV261, Km^r^ | This study |
| pMV261-*cfp-10*^R57A^ (Mm) | *cfp-10^R57A^* (Mm) under the control of the promoter in plasmid pMV261, Km^r^ | This study |
| pMV261-*cfp-10* (Mtb) | *cfp-10* (Mtb) under the control of the promoter in plasmid pMV261, Km^r^ | This study |
| pMV261-*cfp-10*^R57A^ (Mtb) | *cfp-10^R57A^* (Mtb) under the control of the promoter in plasmid pMV261, Km^r^ | This study |
| pGEX6P-1 | *E. coli* GST fusion expression vector with tac promoter, Amp^r^ | Novagen |
| pGEX6P-1-*cfp-10* (Mm) | pGEX6P-1 carrying *cfp-10* (Mm) coding region, Amp^r^ | This study |
| pGEX6P-1*-cfp-10*^R57A^ (Mm) | pGEX6P-1 carrying *cfp-10^R57A^* (Mm) coding region, Amp^r^ | This study |
| pGEX6P-1-*cfp-10* (Mtb) | pGEX6P-1 carrying *cfp-10* (Mtb) coding region, Amp^r^ | This study |
| pGEX6P-1-*cfp-10*^R57A^ (Mtb) | pGEX6P-1 carrying *cfp-10^R57A^* (Mtb) coding region, Amp^r^ | This study |
| pLenti-CMV | Lentiviral transfer vector, human cytomegalovirus immediate-early promoter, Amp^r^ | This study |
| pLenti-CMV-*cfp-10* (Mtb) | pLenti-CMV carrying *cfp-10* (Mtb) coding region, Amp^r^ | This study |
| pLenti-CMV-*cfp-10*^R57A^ (Mtb) | pLenti-CMV carrying *cfp-10*^R57A^ (Mtb) coding region, Amp^r^ | This study |
| psPAX2 | Packaging plasmid psPAX2, encoding HIV-1 gag-pol and rev | This study |
| pMD2.G | Envelope plasmid pMD2.G, encoding VSV-G | This study |
| pCMV-N-*eGFP* | Mammalian expression vector, CMV promoter, N-terminal eGFP-tagged fusion protein for fluorescence imaging | Beyotime |
| pCMV-*cGAS*-*eGFP* | pCMV-N-eGFP carrying *cGAS* coding region, Km^r^ | This study |
| pCMV-*cfp-10* (Mtb) | pCMV carrying *cfp-10* (Mtb) coding region, Km^r^ | This study |
| pCMV-*cfp-10*^R57A^ (Mtb) | pCMV carrying *cfp-10*^R57A^ (Mtb) coding region, Km^r^ | This study |
| pMV261-*esat-6* (Mm) | *esat-6* (Mm) under the control of the promoter in plasmid pMV261, Km^r^ | This study |
| pMV261-*esat-6*^H26A/D30A^ (Mm) | *esat-6^H26A/D30A^* (Mm) under the control of the promoter in plasmid pMV261, Km^r^ | This study |
| pMV261-*esat-6* (Mtb) | *esat-6* (Mtb) under the control of the promoter in plasmid pMV261, Km^r^ | This study |
| pMV261-*esat-6*^H26A/D30A^ (Mtb) | *esat-6*^H26A/D30A^ (Mtb) under the control of the promoter in plasmid pMV261, Km^r^ | This study |
| pET28a | Expression vector with N-terminal His affinity tag, Km^r^ | Novagen |
| pET28a-*esat-6* (Mm) | pET28a carrying *esat-6* (Mm) coding region, Km^r^ | This study |
| pET28a*-esat-6*^H26A/D30A^ (Mm) | pET28a carrying *esat-6*^H26A/D30A^ (Mm) coding region, Km^r^ | This study |
| pLenti-CMV-*esat-6* (Mm) | pLenti-CMV carrying *esat-6* (Mm) coding region, Amp^r^ | This study |
| pLenti-CMV-*esat-6*^H26A/D30A^ (Mm) | pLenti-CMV carrying *esat-6^H26A/D30A^* (Mm) coding region, Amp^r^ | This study |
| pCMV-*esat-6* (Mm) | pCMV carrying *esat-6* (Mm) coding region, Km^r^ | This study |
| pCMV-*esat-6*^H26A/D30A^ (Mm) | pCMV carrying *esat-6^H26A/D30A^* (Mm) coding region, Km^r^ | This study |
| pMV261-Δ*esat-6*Δ*cfp-10* (Mm) | *esat-6* (Mm) and *cfp-10* (Mm) under the control of the promoter in plasmid pMV261, Km^r^ | This study |
| pMV261-*cfp-10-esat-6* (Mtb) | *cfp-10* (Mtb) and *esat-6* (Mtb) under the control of the promoter in plasmid pMV261, Km^r^ | This study |

*Km, kanamycin; Amp, ampicillin; the working concentrations were 50 μg/mL and 100 μg/mL, respectively.

**Table S3. Primers used in this study.**

| **Primers** | | **5′–3′ sequence** | | **Function** |
| --- | --- | --- | --- | --- |
| *cfp-10*-HindIII-F (Mm) | | AAGCTTTCAGAAGCCCATTTGCGAGGACAGCGCCT | | To generate  pMV261-*cfp-10* (Mm) |
| *cfp-10*-BamHI-R (Mm) | | GGATCCATGGCAGAGATGAAGACCGATGC | |  |
| pMV261*-*F | | ATAAAACGAAAGGCCCAGTCTT | |  |
| *cfp-10^R57A^*-BamHI-F (Mm) | | GGATCCTCAGAAGCCCATTTGCGAGGACA | | To generate  pMV261-*cfp-10^R57A^* (Mm) |
| *cfp-10^R57A^*-XbaI-R (Mm) | | TCTAGAATGGCAGAGATGAAGACCGATGCC | |  |
| *cfp-10^R57A^*-F (Mm) | | GCTTCCTGGAA**cgc**GACCACCGC | |  |
| *cfp-10^R57A^*-R (Mm) | | GCGGTGGTC**gcg**TTCCAGGAAGC | |  |
| *cfp-10*-BamHI-F (Mtb) | | GGATCCTCAGAAGCCCATTTGCGAGGACA | | To generate  pMV261-*cfp-10* (Mtb) |
| *cfp-10*-XbaI-R (Mtb) | | TCTAGAATGGCAGAGATGAAGACCGATGCC | |  |
| *cfp-10*-HindIII-F (Mtb) | | AAGCTTTCAGAAGCCCATTTGCGAGGACAGCGCCT | | To generate  pMV261-*cfp-10^R57A^* (Mtb) |
| *cfp-10*-BamHI-R (Mtb) | | GGATCCATGGCAGAGATGAAGACCGATGC | |  |
| *cfp-10^R57A^-*F (Mtb) | | TGCTTCTTGGAA**CGC**CACCACCGCG | |  |
| *cfp-10^R57A^-*R (Mtb) | | CGCGGTGGTG**GCG**TTCCAAGAAGCA | |  |
| *cfp-10*-EcoRI-F (Mm) | | GAATTCATGGCAGAGATGAAGACCGATG | | To generate  pGEX6P-1-*cfp-10* (Mm) |
| *cfp-10*-XhoI-R (Mm) | | CTCGAGTCAGAAGCCCATTTGCGAGGACA | |  |
| *cfp-10^R57A^*-F (Mm) | | GCTTCCTGGAA**cgc**GACCACCGC | | To generate  pGEX6P-1-*cfp-10^R57A^* (Mm) |
| *cfp-10^R57A^*-R (Mm) | | GCGGTGGTC**gcg**TTCCAGGAAGC | |  |
| *cfp-10*-EcoRI-F (Mtb) | | GAATTCATGGCAGAGATGAAGACCGATG | | To generate  pGEX6P-1-*cfp-10* (Mtb) |
| *cfp-10*-XhoI-R (Mtb) | | CTCGAGTCAGAAGCCCATTTGCGAGGACA | |  |
| *cfp-10^R57A^*-F (Mtb) | | CGCGGTGGTG**GCG**TTCCAAGAAGCA | | To generate  pGEX6P-1-*cfp-10^R57A^* (Mtb) |
| *cfp-10^R57A^*-R (Mtb) | | TGCTTCTTGGAA**CGC**CACCACCGCG | |  |
| *cfp-10**attb1*-SpeI-F (Mtb) | | CCTTCTTCTTTTTCCTACAGactagtCCAGTGTGGTGGAATTCTGCAGATATCAacaagtttgtacAAAAAAGCAGGCTTCACCATGGCAGAGATGAAGACCGAT | | To generate  pLenti-CMV-*cfp-10* (Mtb) |
| *cfp-10attb2*-XhoI-R (Mtb) | | GGGGGGGGAGGGAGAGGGGCCTCGAGCGGCCGCCACTGTGCTGGATATCAACCACTTTGTACAAGAAAGCTGGGTTCAGAAGCCCATTTGCGAG | |  |
| *cfp-10^R57A^*-F (Mtb) | | CGCGGTGGTG**GCG**TTCCAAGAAGCA | | To generate  pLenti-CMV-*cfp-10^R57A^* (Mtb) |
| *cfp-10^R57A^*-R (Mtb) | | TGCTTCTTGGAA**CGC**CACCACCGCG | |  |
| *cgas*-HindIII-F | | atgAAGCTTatgcagccttggcacggaaaggc | | To generate  pCMV-*cGAS-eGFP* |
| *cgas*-SalI-R | | atgGTCGACtcaaaattcatcaaaaactggaaa | |  |
| *cfp-10*-HindIII-F (Mtb) | | AAGCTTATGGCAGAGATGAAGACCGATG | | To generate  pCMV-*cfp-10* (Mtb) |
| *cfp-10*-EcoRI-R (Mtb) | | GAATTCTCAGAAGCCCATTTGCGAGGAC | |  |
| *cfp-10^R57A^*-F (Mtb) | | CGCGGTGGTG**GCG**TTCCAAGAAGCA | | To generate  pCMV-*cfp-10^R57A^* (Mtb) |
| *cfp-10^R57A^*-R (Mtb) | | TGCTTCTTGGAA**CGC**CACCACCGCG | |  |
| *ISD*-F | | tacagatctactagtgatctatgactgatctgtacatgatctaca | | To activate cGAS |
| *ISD*-R | | tgtagatcatgtacagatcagtcatagatcactagtagatctgta | |  |
| (Cy3)-*dsDNA100*-F | | ACATCTAGTACATGTCTAGTCAGTATCTAGTGATTATCTAGACATACATCTAGTACATGTCTAGTCAGTATCTAGTGATTATCTAGACATGGACTCATCC | | For confocal microscopy analysis of CFP-10/ESAT-6 (Mtb)and cGAS colocalization |
| *dsDNA100*-R | | GGATGAGTCCATGTCTAGATAATCACTAGATACTGACTAGACATGTACTAGATGTATGTCTAGATAATCACTAGATACTGACTAGACATGTACTAGATGT | |  |
| *esat-6*-BamHI-F (Mm) | | GGATCCATGACAGAACAGCAGTGGAAT | | To generate  pMV261-*esat-6* (Mm) |
| *esat-6*-HindIII-R (Mm) | | AAGCTTTTAAGCAAACATCCCCGTGACGTT | |  |
| pMV261*-*F | | ATAAAACGAAAGGCCCAGTCTT | |  |
| *esat-6^H26A/D30A^*-BamHI-F (Mm) | | GGATCCATGACAGAACAGCAGTGGAAT | | To generate  pMV261-*esat-6^H26A/D30A^* (Mm) |
| *esat-6^H26A/D30A^*-HindIII-R (Mm) | | AAGCTTTTAAGCAAACATCCCCGTGACGTT | |  |
| *esat-6^H26A/D30A^*-F (Mm) | | CCCTC**aGC**CAGAAGGGA**ggc**AATGCTGG | |  |
| *esat-6^H26A/D30A^*-R (Mm) | | CCAGCATT**gcc**TCCCTTCTG**GCt**GAGGG | |  |
| *esat-6*-BamHI-F (Mtb) | | GGATCCctatgcgaacatcccagtgacgtt | | To generate  pMV261-*esat-6* (Mtb) |
| *esat-6*-XbaI-R (Mtb) | | TCTAGAatgacagagcagcagtggaattt | |  |
| *esat-6^H26A/D30A^*-BamHI-F (Mtb) | | GGATCCatgacagagcagcagtggaatt | | To generate  pMV261-*esat-6^H26A/D30A^* (Mtb) |
| *esat-6^H26A/D30A^*-HindIII-R (Mtb) | | AAGCTTctatgcgaacatcccagtgac | |  |
| *esat-6^H26A/D30A^-*F (Mtb) | | cccctc**ggc**aaggaggga**aGC**aatgga | |  |
| *esat-6^H26A/D30A^-*R (Mtb) | | tccatt**GCt**tccctcctt**gcc**gagggg | |  |
| *esat-6*-HindIII-F (Mm) | | acgAAGCTTATGACAGAACAGCAGTGGAATTTC | | To generate  pET28a-*esat-6* (Mm) |
| *esat-6*-NotI-R (Mm) | | atcGCGGCCGCTTAAGCAAACATCCCCGTGACGTT | |  |
| *esat-6^H26A/D30A^*-F (Mm) | | AGCATT**gcc**TCCCTTCTG**GcT**GAGGGCAAG | | To generate  pET28a-*esat-6^H26A/D30A^* (Mm) |
| *esat-6^H26A/D30A^*-R (Mm) | | CTTGCCCTC**AgC**CAGAAGGGA**ggc**AATGCT | |  |
| *esat-6attb1*-SpeI-F (Mm) | | CCTTCTTCTTTTTCCTACAGactagtCCAGTGTGGTGGAATTCTGCAGATATCAacaagtttgtacAAAAAAGCAGGCTTCACCATGACAGAACAGCAGTGGAAT | | To generate  pLenti-CMV-*esat-6* (Mm) |
| *esat-6attb2*-XhoI-R (Mm) | | GGGGGGGGAGGGAGAGGGGCctcgagCGGCCGCCACTGTGCTGGATATCAaccactttgtacaagaaaGCTGGGTTTAAGCAAACATCCCCGTG | |  |
| *esat-6^H26A/D30A^*-F (Mm) | | AGCATT**gcc**TCCCTTCTG**GcT**GAGGGCAAG | | To generate  pLenti-CMV-*esat-6^H26A/D30A^* (Mm) |
| *esat-6^H26A/D30A^*-R (Mm) | | CTTGCCCTC**AgC**CAGAAGGGA**ggc**AATGCT | |  |
| *esat-6*-EcoRI-F (Mm) | | GAATTCATGACAGAACAGCAGTGGAAT | | To generate  pCMV-*esat-6* (Mm) |
| *esat-6*-XhoI-R (Mm) | | CTCGAGTTAAGCAAACATCCCCGTGACG | |  |
| *esat-6^H26A/D30A^*-F (Mm) | | AGCATT**gcc**TCCCTTCTG**GcT**GAGGGCAAG | | To generate  pCMV-*esat-6^H26A/D30A^* (Mm) |
| *esat-6^H26A/D30A^*-R (Mm) | | CTTGCCCTC**AgC**CAGAAGGGA**ggc**AATGCT | |  |
| *cfp-10*-EcoRI-F (Mm) | atGAATTCATGGCAGAGATGAAGACCGATG | | | To generate  pMV261-*esat-6cfp-10* (Mm) |
| *cfp-10*-R (Mm) | TTCCACTGCTGTTCTGTCATGAAGCCCATTTGCGAGGACA | | |  |
| *esat-6-*F (Mm) | TGTCCTCGCAAATGGGCTTCATGACAGAACAGCAGTGGAA | | |  |
| *esat-6*-HindIII-R (Mm) | taAAGCTTTTAAGCAAACATCCCCGTGACGT | | |  |
| pMV261*-*F | ATAAAACGAAAGGCCCAGTCTT | | |  |
| *cfp-10^R57A^b*-EcoRI-F (Mtb) | GAATTCATGGCAGAGATGAAGACCGATGC | | | To generate  pMV261-*cfp-10-esat-6* (Mtb) |
| *cfp-10^R57A^b*-R (Mtb) | tccactgctgctctgtcatTCAGAAGCCCATTTGCGAGGA | | |  |
| *esat-6^H26A/D30A^b*-F (Mtb) | TCCTCGCAAATGGGCTTCTGAatgacagagcagcagtgga | | |  |
| *esat-6^H26A/D30A^b*-HindIII-R (Mtb) | AAGCTTctatgcgaacatcccagtgac | | |  |
| *actb*-F | | | CATTGCTGACAGGATGCAGAAGG | For RT-qPCR (for mouse) |
| *actb*-R | | | TGCTGGAAGGTGGACAGTGAGG |  |
| *ifnb1*-F | | | GCCTTTGCCATCCAAGAGATGC |  |
| *ifnb1*-R | | | ACACTGTCTGCTGGTGGAGTTC |  |
| *ifit1*-F | | | TACAGGCTGGAGTGTGCTGAGA |  |
| *ifit1*-R | | | CTCCACTTTCAGAGCCTTCGCA |  |
| *ifit2*-F | | | CGAACTACCGTCTGGATGACTG |  |
| *ifit2*-R | | | CTTCAACCAGCGCCATTGCTTG |  |
| *ifit3*-F | | | GCTCAGGCTTACGTTGACAAGG |  |
| *ifit3*-R | | | CTTTAGGCGTGTCCATCCTTCC |  |
| *irf7*-F | | | CCTCTGCTTTCTAGTGATGCCG |  |
| *irf7*-R | | | CGTAAACACGGTCTTGCTCCTG |  |
| *isg15*-F | | | CATCCTGGTGAGGAACGAAAGG |  |
| *isg15*-R | | | CTCAGCCAGAACTGGTCTTCGT |  |
| *cxcl10*-F | | | ATCATCCCTGCGAGCCTATCCT |  |
| *cxcl10*-R | | | GACCTTTTTTGGCTAAACGCTTTC |  |
| *tnfa*-F | | | GGTGCCTATGTCTCAGCCTCTT |  |
| *tnfa*-R | | | GCCATAGAACTGATGAGAGGGAG |  |
| *il1b*-F | | | TGGACCTTCCAGGATGAGGACA |  |
| *il1b*-R | | | GTTCATCTCGGAGCCTGTAGTG |  |
| *Ifng*-F | | | ATGAACGCTACACACTGCATC |  |
| *Ifng*-R | | | CCATCCTTTTGCCAGTTCCTC |  |
| *Ifna2*-F | | | TACTCAGCAGACCTTGAACCT |  |
| *Ifna2*-R | | | CAGTCTTGGCAGCAAGTTGAC |  |
| *actbZ*-F | | | CGAGCAGGAGATGGGAACC | For RT-qPCR (for zebrafish) |
| *actbZ*-R | | | CAACGGAAACGCTCATTGC |  |
| *ifnphi1Z*-F | | | GTCAAAGGTGGAGGACCAGG |  |
| *ifnphi1Z*-R | | | CCCTGGCATCCATAAGGTGG |  |
| *isg15Z*-F | | | CCAACGAGACCGTCGATCAG |  |
| *isg15Z*-R | | | CTCGAGCTGTCTGCCTTTGA |  |
| *tlr7Z*-F | | | TTTCAAAGCCTGGCCGATCT |  |
| *tlr7Z*-R | | | CTGGTGCACGTGATCTCCTT |  |
| *nod2Z*-F | | | CTGAGGAGCAAAAGGCTGGA |  |
| *nod2Z*-R | | | GTTGAAGTTGCAGTGGCTGG |  |
| *ripk2Z*-F | | | ATATCAGACAGCGCAGGTCG |  |
| *ripk2Z*-R | | | AGGGGTTGTATGGCAAGAGC |  |
| *relaZ*-F | | | GCATCCCTGGAGAGAAGAGC |  |
| *relaZ*-R | | | AGGAGATCCGCACTCTGACT |  |
| *mxcZ*-F | | | GAGGACCTGGAAGAACCTGC |  |
| *mxcZ*-R | | | GAGCCCACAGAGGAGTGAAC |  |

*The underlined sequences indicate the restriction endonuclease recognition sites, and the bolded nucleotides represent the mutation sites.
